# Systematic mapping of neural signals in the periphery reveals a rich and specific alphabet for immune cell communication

**DOI:** 10.1101/2022.12.29.522179

**Authors:** Karen Regev Berman, Neta Milman, Meital Segev, Elina Stratovsky, Shai S. Shen-Orr

## Abstract

Accumulating evidence indicate a strong link between neural signals and the immune system. Given neural signals constitute a large family that may be leveraged for communication, we systematically explored the neuro-immune regulation network in the periphery and uncovered a broad yet lineage selective expression of neuro-receptors on immune cells. We constructed a rich social immune network map showing the neural molecular pathways supporting the regulation of the immune system at steady state. Our results emphasize neuro-receptors role in the commitment and differentiation of B and T cells along their developmental process. We identified the immune cells’ functionality in the specific tissue is extensively shaped by the communication with the microenvironment and nervous systems via a rich alphabet of neural mediators. Collectively, our findings suggest neural genes are an integral part of the immune regulatory system and provide clear testable new avenues of experimental follow up for neuroimmunologists and immunologists alike.

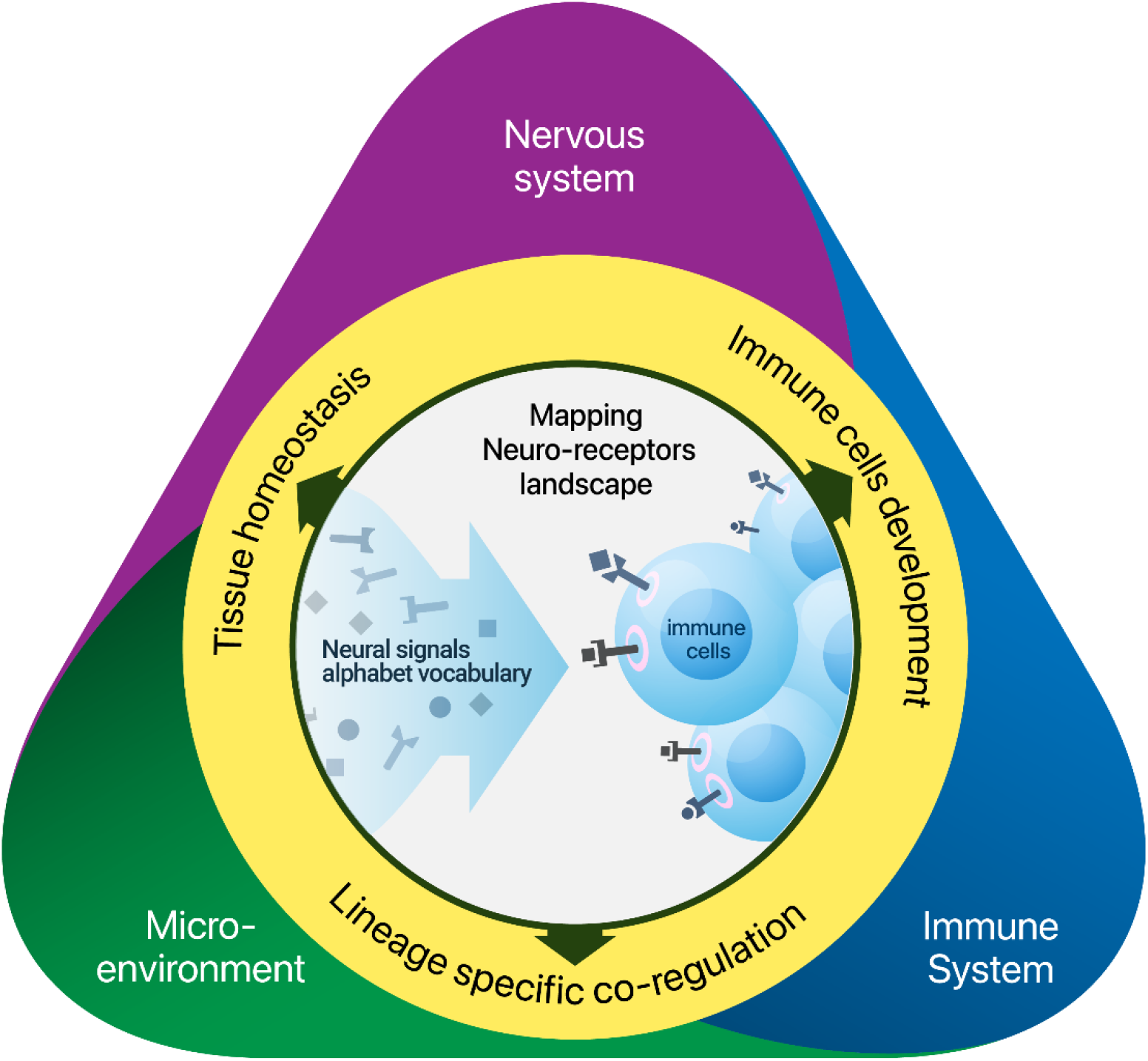

**Highlights:**

- 160 neural genes potentially active in immune cells in the periphery with high specificity to lineage
- Neuro-receptors co-expressed in immune lineages are enriched for biological functions
- Changes in neuro-receptors expression influence the HCS differentiation and commitment
- Immune-microenvironment neural signaling architecture reflects tissue biological role

## Introduction

Studies in the growing field of neuroimmunology established that immune processes in peripheral organs are also modulated by neural signals. Neurotransmitters, neuropeptides and neurohormones exert their effect by binding to their respective receptors expressed on immune cells^1^, including hallmark regulation by norepinephrine, acetylcholine, dopamine and more^1–6^. However, given the large number of different neural signals and the many immune cell-types and the historical obliviousness of mainstream immunology research to neural regulation of immune cells, it is likely that only a small fraction of neural-immune interactions has been reported. The diversity of neural signaling genes expressed in different immune cell types may thus underlie an entire uncharted alphabet of neuro-immune communication.

Previous work by us and others established a global, immune-centric view of the inter-cellular communication via cytokines and chemokines (hereon cytokines)^7–9^. The systematic mapping of the immune cell-cytokine milieu enables the discovery of novel interactions, and the prediction of previously unreported associations of cytokines with disease conditions. These studies highlight that the diversity of receptors on immune cells is key for understanding of how the immune system responds and adapts to perturbations. Notably, these studies have predominantly focused on cytokine signaling due to their perception as the main players of immune cell communication, overlooking other receptors on immune cells. The accumulating evidence for the neuro-immune interactions raises the possibility that neural signaling may have been systematically leveraged either for brain-immune communication or even co-opted for communication between immune cells themselves or their microenvironment. Mapping the social network of immune cells using the neural signaling alphabet may thus dramatically broaden our understanding of immune function and its integration with other systems.

Here we systematically mapped the neural-immune communication potential by studying the expression of hundreds of neural signals and receptors across a broad range of immune cells in steady state conditions. For this, we analyzed The Immunological Genome Project (ImmGen)^10^, a resource spanning whole genome gene expression profiles of mouse immune cells across all lymphoid organs and other tissues which are monitored by immune cells. We mapped the landscape of neuro-receptor expression on immune cells and detected a broad range of neural signaling genes invoking inter-cellular regulatory structures. We constructed communication networks between cells via neural signals, allowing to decipher the architecture of neural signal communication between immune cells and their microenvironment and nervous system in a tissue dependent manner. Lastly, we examined co-expression relations between neuro-receptors and their shared function in immune processes, with specific emphasis on the effect of neural signals on hematopoiesis, which we also validated experimentally.

## Results

To explore the systemic effect of neural signals on the immune system in the periphery, we examined the expression of neural signaling receptors in immune cells. We focused on 257 receptors of neurotransmitters, neuropeptides and neurohormones, and their respective 61 ligands (See Methods for gene selection and Table S2 for gene list). We analyzed whole genome expression data derived from mouse sorted cells obtained from ImmGen, a large high-quality database characterizing murine immune cells^10^. Particularly, we analyzed 180 immune cell types across the entire murine hematopoietic lineage from 19 lymphoid organs and other tissues in the periphery in steady state conditions [Table S1, Figure S1].

### Neuro-receptor expression profile distinguishes between immune lineages

To explore the global landscape of immune potential response to neural signaling, we projected the 257 neuro-receptors expression profile of each immune cell to a low-dimensional representation using principal component analysis (PCA). PCA showed a broad organization of immune cells, with the first two principal components explaining a total of 35% of variation in neuro-receptors expression [Figure 1A]. Comparable percent of variance (42%) was explained by the collection of all cytokine receptors in a similar PCA analysis [Figure S2E]. The organization of the immune cells appeared to be driven by immune cell lineage, but not by tissue of origin [Figure S2A]. These results indicate that the expression of neuro-receptors is distinctly different on cells from different lineages. Moreover, this PCA spread showed striking significant separation between progenitor cells, lymphoid and myeloid cell types (Figure 1B top row, p< 2.05E-05, see Methods for separation measure), suggesting neuro-receptors expression on immune cells follow a pre-determined developmental program. To evaluate if immune cell lineage separation is a property of expression specific to neuro-receptors we compared it against a random set of genes. We observed that the expression of neuro-receptors separates lymphocyte cells from myeloid and progenitor cells in a significant manner (p <0.0001, Figure S2B, not observed for progenitor-myeloid cell separation). Only a partial set of canonical neural signals have been studied in peripheral signaling in the immune system among them are acetylcholine, norepinephrine, dopamine, neuromedin U, vasoactive intestinal peptide, substance P^3^. Restricting immune-cell expression profile only to these canonical neural signaling receptors (a total of 41 genes, Figure S2C), we observed a weaker separation in PCA in terms of distance between groups of lymphocytes, myeloid and progenitor cells than that observed for the entire neuro-receptor expression profiles (Figure 1B bottom row). This could not be explained by the potential confounder of difference in gene set size (Figure S2D, pvalue < 1.2E-7 for comparing separation between groups). Together, these results suggest a much broader neural alphabet is in use than appreciated to date and this alphabet is closely involved with multiple immune cell types.

**Figure 1:**
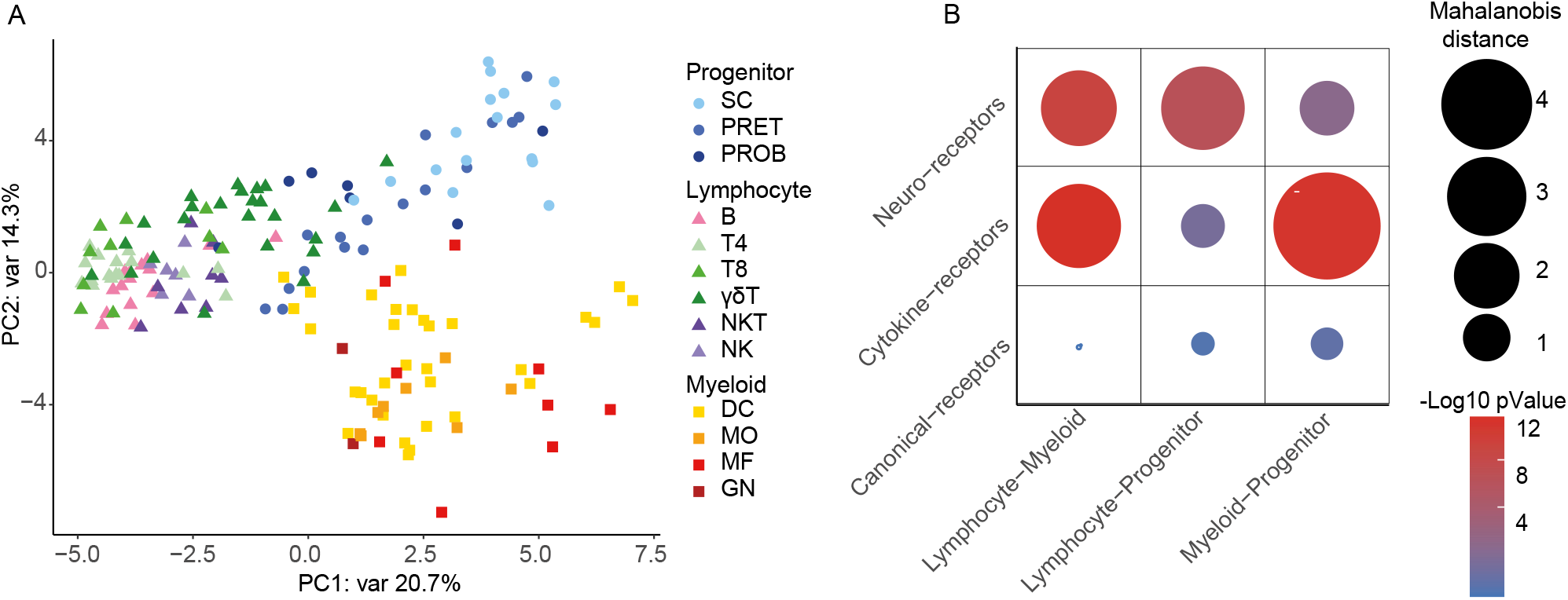
Neuro-receptor expression profile distinguishes between immune lineages. **(A)** PCA based on neuro-receptors log2 expression for all immune cell types (mean of replicates) shows seperation of cells by their lineage and their lineage type (lymphocyte, meyloid or progenitor cell) indicated by color and shape respectively. Immune lineages included: stem cells (SC), progenitor B cells (PROB), progenitor T cells (PRET), B cells (B), CD4 T cells (T4), CD8 T cells (T8), gamma-delta T cells (ɣδT), natural killer T cells (NKT), natural killer (NK), dendritic cells (DC), monocytes (MO), macrophages (MF), and granulocytes (GN). **(B)** Heatmap depicting the multivariate distance in PCA space between lymphocytes, meyloids and progenitor cell subsets based on either neuro-receptors, canonical receptors or cytokine receptors expression. Circle size represents Mahalanobis distance between groups, color represents the -log10 pValue for separation significance tested by t-test.

To appreciate neural signals importance as regulators of the immune system we quantified neuro-receptors organization on immune cells compared to cytokine receptors, a well-established family of regulators of the immune system. We repeated the PCA with immune cells now described by their cytokine receptors expression profile (a total of 110 cytokine receptors, Figure S2E, Table S4) and analyzed for differences in separation of immune cell lineages. We observed a higher separation between lymphocytes and progenitor cells based on neuro-receptor profile than by cytokine receptor profile (Figure 1B middle row, Figure S2F, pvalue < 2.3E-10 when controlling for group size). This was primarily due to B-cells, whose neural receptor profile was similar to other lymphocytes but distinct when analyzed based on cytokine receptor profile. These neuro-receptor lineage specific profile raise the possibility neuro-receptors adopted specific functionality in the immune system, with distinct roles in the development process, the adaptive and innate systems.

### Neuro-receptors are broadly yet selectively expressed on immune cells

The gene expression profile of immune cells provides quantitative information to approximate a global picture of inter-cellular signaling^11^. To map the roles neural signaling play in immune regulation requires knowledge of their cell type activity and specificity. Estimating genes functionality from expression data requires distinguishing between functional expression and noise level. Despite clear differences in dynamic ranges of different genes, a common practice when analyzing microarray expression data is to set a global threshold across all genes to what would be considered functional expression. Setting a global expression threshold [Figure S3A], akin to ImmGen consortium^12^, we observed that 68% of neural genes (receptors and ligands) expression values lie in an intermediate expression range for which it is difficult to determine with high probability if truly expressed or not [Figure 2A]. Furthermore, we observed a non-uniform distribution of neural genes expression among cell-types, with samples below threshold biased towards association with a particular cell-lineage. For example, we noted Hrh4 histamine receptor as selectively expressed in natural killer T (NKT) cells in concordance with its role to ensure optimal IL-4- and IFN-δ production by NKT cells^13^ [Figures 2B]. Yet we noted that Hrh4’s expression values were below the global expression threshold and would thus be considered not expressed [Figures 2A]. This non-random, low, selective expression signal motivated us to develop a gene-specific expression threshold algorithm (see Methods, [Figure S3C]). Applying this gene specific threshold allowed us to rescue 12 neural signals and 72 neuro-receptors selectively expressed in immune cells [Figure 2C]. These expression values would be considered undetectable using the global threshold. Together, these two approaches yielded a total of 160 neural genes (48 ligands and 122 receptors) potentially active in the immune system [Figure 2C]. We observed high specificity to lineage with 27.5% of the neuro-receptors expressed in a single lineage, and 12.5% were expressed in all lineages functioning as housekeeping genes [Figure 2D, Table S2]. In total, the immune system has the potential to respond to 75% of the neural signals through their corresponding neuro-receptors [Figure 2E]. Taken together, we detected that the immune system expresses a broad range of neural signaling genes suggesting a broader involvement of neural signals in immune communication than appreciated to date.

**Figure 2:**
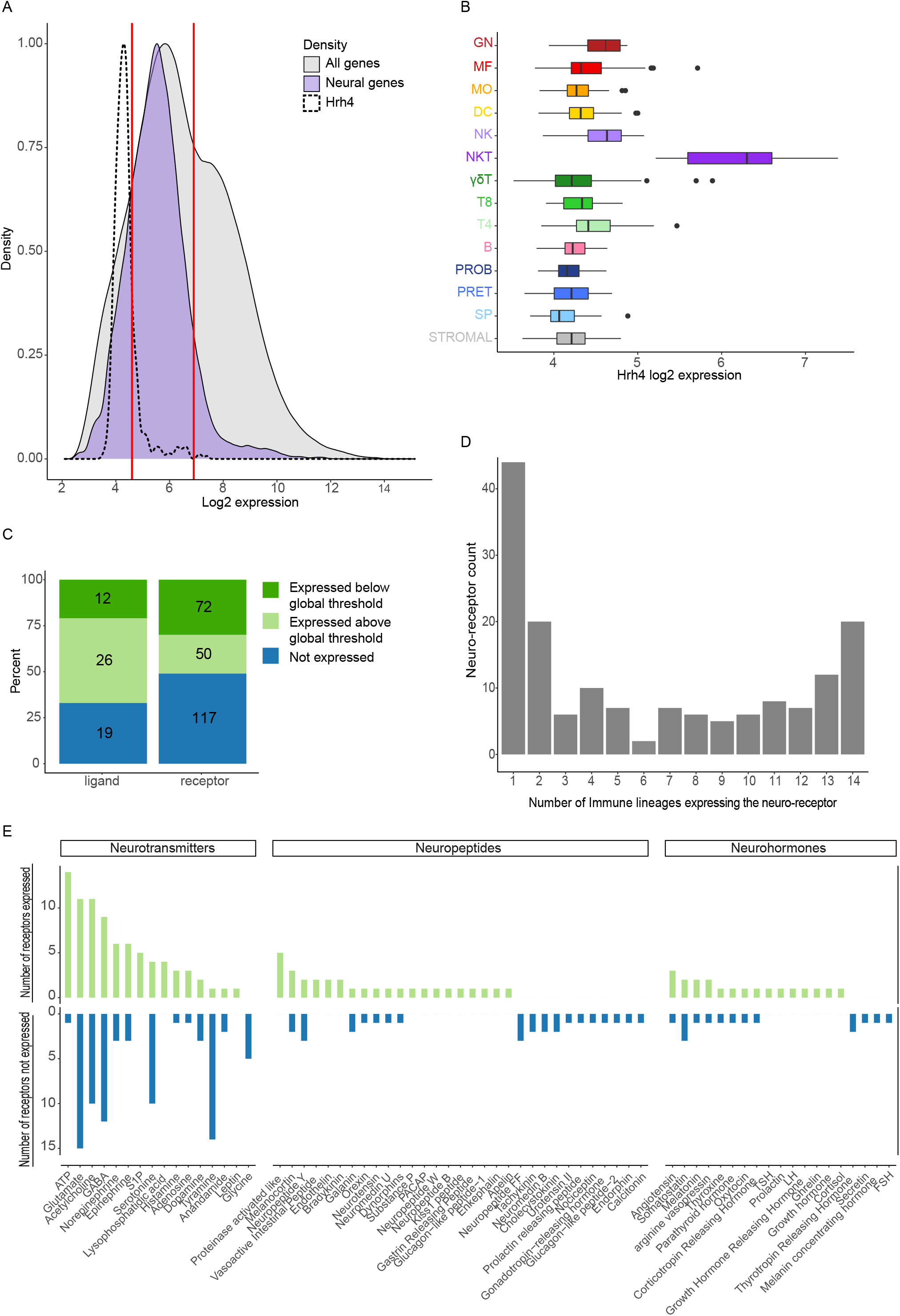
Neuro-receptors are broadly yet selectively expressed on immune cells. **(A)** A need for gene specific threshold for determining expression. Log2 expression distribution of all genes (light grey) and of neural genes only (purple) across 567 samples of immune cells in ImmGen dataset. For reference, overlaid is the distribution of the histamine receptor gene Hrh4 across all cell subsets (dotted line) with its expression in natural killer T cells (NKT) marked. The red line at log2 expression of 6.9 indicates the global threshold of expression and the red line at log2 expression of 4.6 indicates the threshold of background signal (see Figure S3A). Between these two lines is the intermediate expression range for which it is difficult to determine with high probability if truly expressed or not. **(B)** Box plot of histamine receptor gene Hrh4 log2 expression values across 567 samples of immune cells separated by lineage. Bar color and y axis labels indicate immune lineage. Hrh4 log2 expression below global threshold of 6.9 is biased towards association with a particular cell-lineage, NKT. Immune lineages included: stem cells (SC), progenitor B cells (PROB), progenitor T cells (PRET), B cells (B), CD4 T cells (T4), CD8 T cells (T8), gamma-delta T cells (ɣδT), natural killer T cells (NKT), natural killer (NK), dendritic cells (DC), monocytes (MO), macrophages (MF), and granulocytes (GN). **(C)** Cumulative bar plot of the expression statistics for neuro-receptors and neural ligands. The number of genes indicated on the bars represent:genes that did not pass gene specific threshold and are not expressed in immune cells (blue), genes expressed in immune cells with expression values above global threshold (light green) and ‘rescued’ genes expressed in immune cells with expression values below global threshold of 6.9. (**D**) Bar plot summarizing the number of expressed neural genes by the number of lineages the gene is expressed in. (**E**) Bar plot indicating for each neural signal the number of neuro-receptors it is known to bind that are expressed in immune cells (top, green) and the number of neuro-receptors it is known to bind that are not expressed in immune cells (bottom, blue). Neural signals groups are separated by type: neurotransmitter, neuropeptide or neurohormone, and ordered by the number of expressed neuro-receptors within each group.

### Neural signals create tissue specific neuro-immune-stromal networks

Our systems-wide analysis of neural genes expression in the periphery strongly supports that the effect of neural signals on the immune system is comprehensive. To build an understanding of how cells communicate through these signals we mapped the source of neural signaling and identified interactions between cells through the expression of neural ligands and their receptors in a tissue dependent manner. We focused on the gastrointestinal (GI) tissue (see Methods), where the neuro-immune axis is known to regulate gastrointestinal barrier function and host protection^14^. We mapped a neuro-immune communication network in the intestine, analyzing neuro-receptors and ligands of immune cells from ImmGen data and neural ligands of the enteric neurons from single cell data set^15^ [Figure 3A]. The communication network showed that neurons in the intestine mainly interact with ɣδ T cells, known to comprise approximately 50% of total T cells in the small intestine^16^. ɣδ T cells’ role in promoting tissue repair and homeostasis^17^ is compatible with their intercellular communication through neural signals known to act as growth factors or growth factors stimulators such as glucagon-like peptide-1, enkephalin, galanin, ghrelin, vasoactive intestinal peptide. Moreover, looking at the intestine network highlight that immune cell regulation via neural signals may act in either autocrine or paracrine manner. For example, GABA, known to down-regulate pro-inflammatory cytokines^16^, is potentially involved both in ɣδ T cells autocrine regulation as well as in regulating dendritic cells which express GABA receptors. Of note, we did not detect neuronal expression of GABA by enteric neurons in the tissue [Figure 3A]. Macrophage function in the intestine differs by location with lamina propria macrophages exerting pro-inflammatory functions and muscularis macrophages exerting tissue protective functions^18^. We observed macrophage dependent adrenergic receptor subtype expression compatible with their function. Specifically, β_2_ adrenergic receptor, known to exert anti-inflammatory effects^18^, is highly expressed on tissue-protective muscularis macrophages whereas β1 adrenergic receptor, known to exert pro-inflammatory effects^19^, is expressed on lamina propria macrophages which are highly phagocytic [Table S5]. We also observed that lamina propria macrophages are regulated by nitrergic enteric neurons through vasoactive intestinal peptide potentially suppressing their inflammatory response and differentiation^20^.

**Figure 3:**
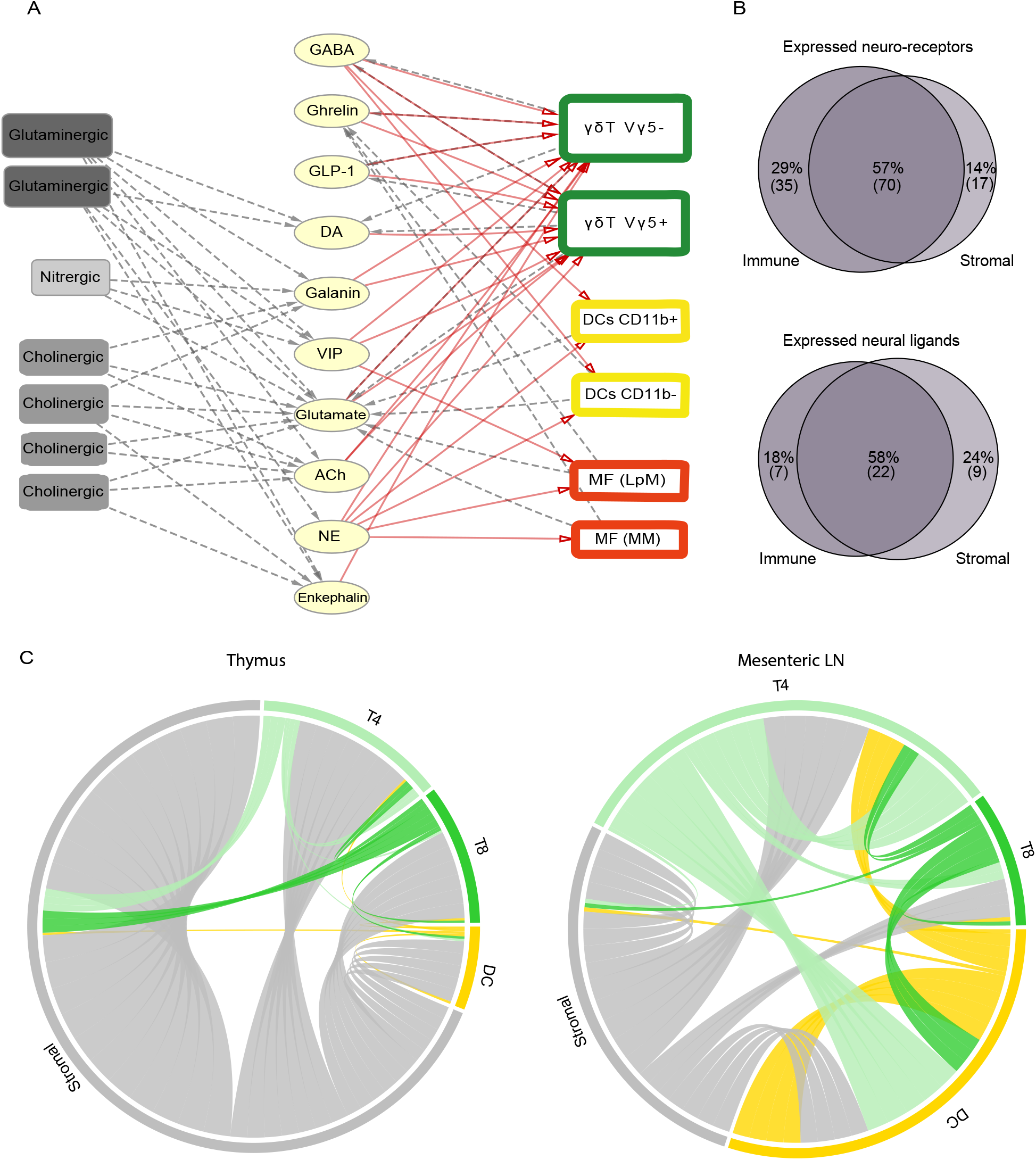
Neural signals create tissue specific neuro-immune-stromal networks. Cell-cell communication network by neural signals at the tissue level. **(A)** Network of neuro-immune potential interactions in the intestine. Communication between neurons (left) and immune cells (right) through specific neural signals (middle) where the cell expressing the ligand is connected by grey dotted edge with direction from the cell to the neural signal, and cell expressing the neuro-receptor is connected by red edge with direction from the neural signal to the cell, giving a complete direction of the potential interaction. Lineages included: gamma-delta T cells (ɣδT) Vγ5+ TCRs and Vγ5-TCRs populations, dendritic cells (DC) CD103+CD11b+ and CD103+CD11b-populations, and macrophages (MF) residing in the lamina propria (LpM) and muscularis (MM) compartments. GLP1-glucagon-like peptide-1, DA-dopamine, VIP-vasoactive intestinal peptide, Ach-achetylcholine, NE-neuropenipherine. For the specific neuro-receptor expressed on immune cells see Table S5. **(B)** Venn diagrams describing the number and percent of expressed receptors (top) and ligands (bottom) shared by the immune cells and stromal cells. Note that stromal cells’ expression profile was evaluted from 3 tissues compared to 19 tissues of immune cells. **(C)** Circus plots comparing stromal-immune inter-cellular neural signaling network in the thymus (left) and mesenteric lymph node (LN) (right) reflecting tissue specific regulation of T cell differentiation and activation. The communication network includs cell types represented in both tissues: T8 cells, T4 cells, DC and epithelial stromal cell. Connections are made between a cell expressing neural ligand with cell expression the respective neuro-receptor. Edge color indicates the lineage of the cell expressing the ligand in the potential interaction.

Immune cell function, activation and migration is known to be regulated also by the microenvironment, though it has predominantly been ascribed to the regulation via cytokines and chemokines^21,22^. We tested the extent to which the microenvironment leverages neural signals for communication with the immune cells. We noted that 57% of the neuro-receptors and 58% of the ligands are expressed by both immune and stromal cells in the different tissues in ImmGen data [Figure 3B]. The high percent of shared neuro-receptors and ligands suggests a tri-directional use of the neural genes in regulating neuro-immune-microenvironment interactions. We thus extended our mapping of neural signal communication to include the interactions of stromal and immune cells and observed that this inter-cellular network signaling architecture can vary between tissues reflecting the tissue biological role. For example, in T cell differentiation and activation we observed that in the thymus the neural signals predominantly originated from stromal cells, providing ques essential for thymocyte development^23^, whereas in the mesenteric lymph node T cell receive greater extent of neural signals from immune cells [Figure 3C]. Specifically, we observed the emergence of DC signaling, presumably reflecting DC role during priming of CD4+ T cells^24^. Thus, the function of immune cells in the tissue is extensively shaped by a rich alphabet of neural signals whose communication with immune cells is tissue specific.

### Neuro-receptors immune cell expression shows selective pressure for within-lineage variability

To ensure proper immune response, immune cells are tightly controlled during development and in response to internal or external stimuli, which modifies the functional and structural roles of the cells. To capture the differences in the landscape of immune cells responses to neural signaling we built on concepts defined for studying the complexity of promoter regulation and transcriptome dynamics^25,26^. We calculated for each cell-type a diversity index, telling of the breadth of neuro-receptors “repertoire” and a specialization index, indicative of how unique the neuro-receptors expressed on the cell-type compared to other cells (see Methods, Figure 4A, Table S1). We detected a negative association between the immune cell diversity and specialization (Figure 4B, r =-0.57 by Pearson’s, p-value<0.0001) implying that cells expressing highly specific neuro-receptors respond to a narrow range of neural signal stimuli, whereas cells with higher diversity correspond to a more uniform distribution of neuro-receptor expression frequencies. Moreover, we observed this negative association of neuro-receptor specialization and diversity independently in all immune cell-lineages but monocytes (Figure 4C, Figure S4B). This strong negative correlation was not observed for cytokine receptors, neither for the overall global correlation nor the lineage specific one (Figure S4A, r=-0.19 for global, Figure 4C for lineage specific contrasts). The negative association of specialization and diversity within lineage appears to build up on the importance of neuro-receptors in guiding lineage decisions; We noted that the stem cells centroid has the highest specialization score whereas the PreT and ProB progenitor centroids have the highest diversity scores [Figure 4B]. This suggests that at early stages of immune cells development the neural regulation is highly specific, whereas at later stages of development the repertoire of neuro-receptors expressed changes.

**Figure 4:**
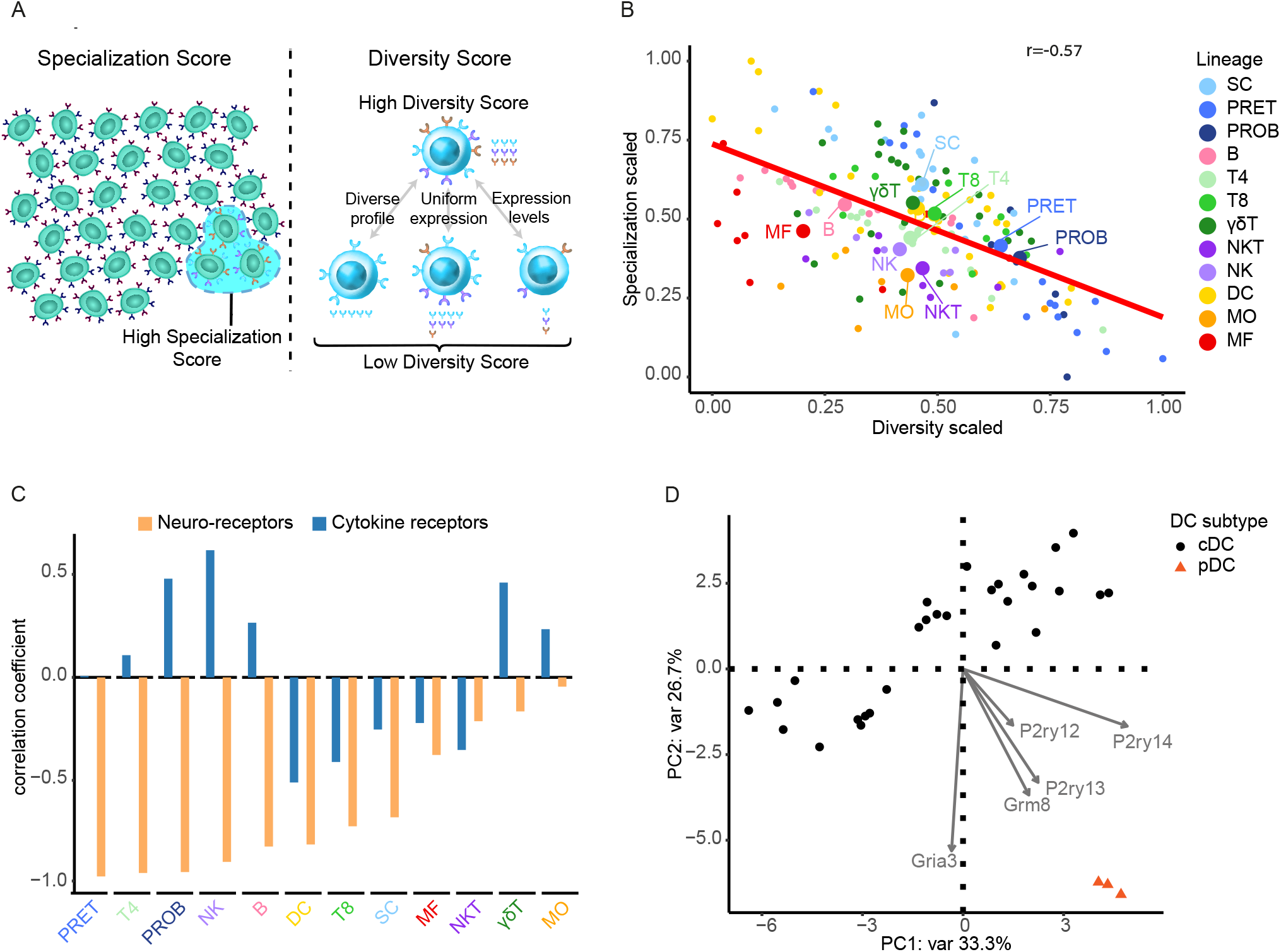
A negative correlation between cell potential to respond to neural signals and the specificity of signals it can respond to. **(A)** Illustrative discription of cells specialization and diversity scores. We calculated for each cell-type a diversity index, telling of the breadth of neuro-receptors “repertoire” and a specialization index, indicative of how unique the neuro-receptors expressed on the cell-type compared to other cells. **(B)** Scatter plot of cell diversity versus specialization scores. Scores are scaled ranging 0 (low) to 1 (high). A negative correlation is observed between diversity and specialization of immune cell types (red line, Pearson’s correlation coefficient r= -0.57). Cells colored by lineage, and lineage centroids are enlarged and labeled. Immune lineages included: stem cells (SC), progenitor B cells (PROB), progenitor T cells (PRET), B cells (B), CD4 T cells (T4), CD8 T cells (T8), gamma-delta T cells (ɣδT), natural killer T cells (NKT), natural killer (NK), dendritic cells (DC), monocytes (MO) and macrophages (MF). **(C)** Bar plot of the linear regression correlation coefficient of diversity vs. specialization calculated for each lineage cell types with cell type profiles defined either solely by neuro-receptor (orange) or by cytokine receptors (blue). Negative correlation between diversity and specialization is more dominant in neuro-receptors than in cytokine receptors. **(D)** PCA of DCs based on expression profile of 122 expressed neuro-receptors separates plasmacytoid DC (pDC) from conventional DC (cDC). Neuro-receptors with loading score < -0.15 in PC2 are marked with arrow showing the loading.

The observation that many neuro-receptors expression profiles are lineage specific [Figure 2D], in combination with the negative association of specialization and diversity we noted within lineage, suggests that neuro-receptors are likely to be involved in regulation of lineage specific states or processes. For example, we observed that plasmacytoid dendritic cells (pDC), have the highest specialization scores and very low diversity scores [Table S1]. This indicates pDCs are specialized cells in the context of neural communication expressing highly specific neuro-receptors. Performing PCA for DCs based on their neuro-receptors expression profile, grouped pDCs separately based on the glutamate receptors Grm8, Gria3 and the purinergic receptors (P2Y) [Figure 4D, Figures S4C]. P2Y signaling in pDCs up-regulates chemokine receptor CCR7 and therefore pDCs undergo phenotypic maturation and migration^27^. Glutamate receptors showed strong pDC association, possibly extending the known role of the glutamate receptor family as mediator of DC role in driving a regulatory T cell response^28^. Thus, our analysis highlights that neural signals may have an impact on the development of the immune system and identifies putative novel neuro-receptors associations.

### Neuro receptors are co-expressed in a lineage specific manner

Receptor co-expression is a fundamental mechanism by which immune cells ensure proper functionality and its detection has historically helped advance deciphering complex processes such as multi-step activation, immune checkpoints, and cytokine signaling^29,30^. Thus, we investigated which neuro-receptors have similar expression patterns across the different immune lineages and therefore might be involved in the same immune regulatory pathway. We identified nine co-expression modules of highly correlated neuro-receptors which are significantly associated with one or more lineages [Figure 5A, Figure S5, Table S6]. Enrichment analysis revealed specific biological functions for modules associated with expression in stem cells, myeloid cells, and B cells but not in other cells of the adaptive system [Figure 5C, Table S6]. As relatively little is known about neuro-receptors signaling function in immune cells, we reasoned that studying neuro-receptors co-regulation with their better-known cytokine counterparts will aid in understanding their function in immune lineages. We clustered neuro-receptors and cytokine receptors co-expressed in specific lineages [Figure 5B, Table S7] which yielded broader insight on biological functions [Figure 5C, Table S8].

**Figure 5:**
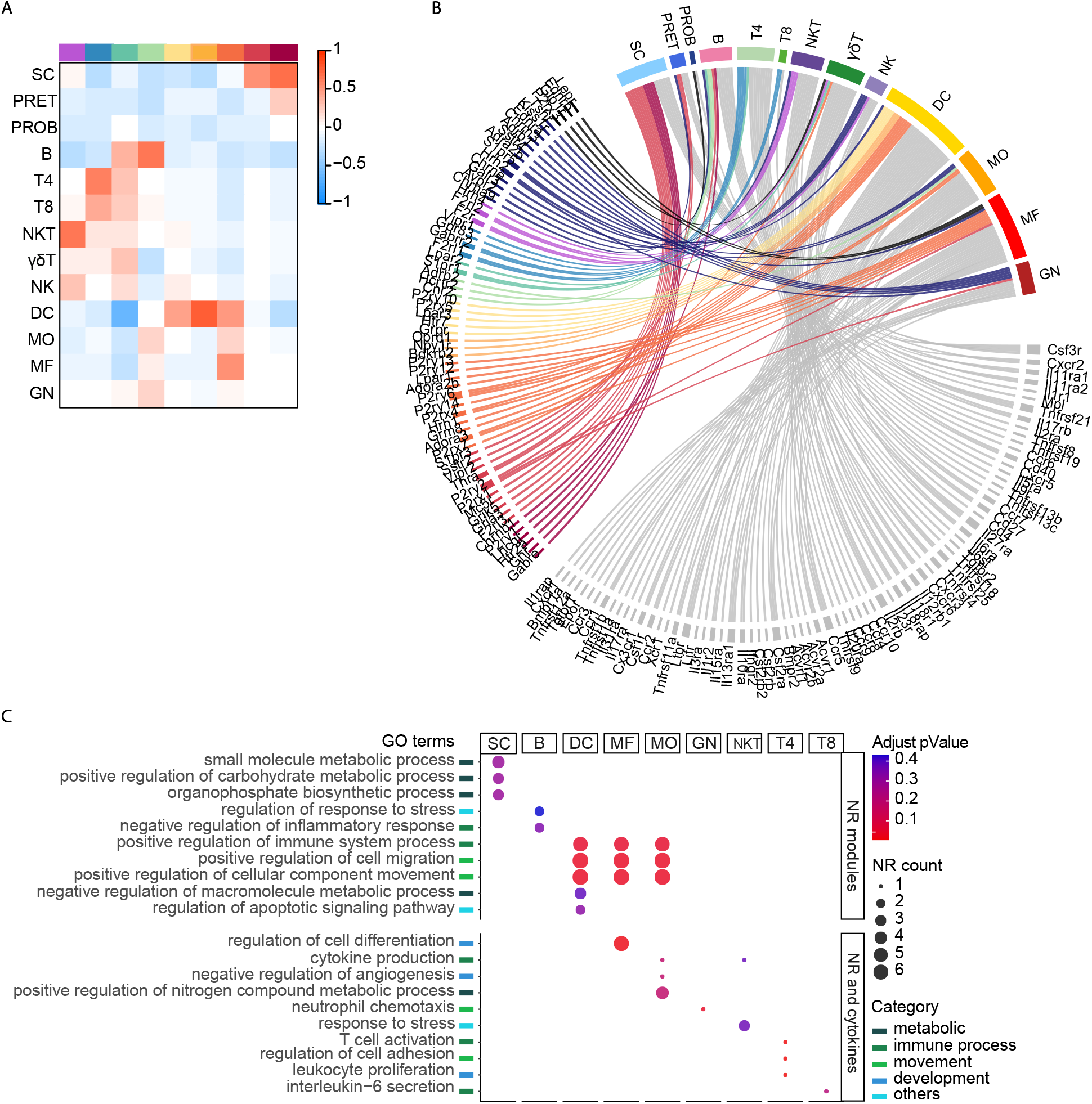
Neuro receptors are co-expressed in a lineage specific manner. 122 expressed neuro-receptors were grouped into modules based on co-expression analysis. The modules genes expression profile across immune cell types significantly correlated to one or more lineages. (**A**) Heatmap depicts the module-lineage (columns and rows respectively) association by correlation (color). (**B**) Neuro-receptor / cytokine receptor – lineage association. Circus plot representing association between neuro-receptor / cytokine receptor with expression profile across immune cell types significantly correlated with the specific lineage (Bonferroni corrected *p-value* < 0.05). Interactions are colored by neuro-receptor module or cytokine receptor (grey). (**C**) Enriched GO terms in neuro-receptors co-expressed with lineage specific profile. Top: For neuro-receptor gene sets identified by co-expression modules. Bottom: For combined list of neuro- and cytokine receptors significantly correlated to lineage. Color indicates Benjamini-Hochberg adjusted enrichment *p-value*. Size indicates the number of neuro-receptors enriched in a GO term. Go terms are annotated by catergory. Immune lineages: stem cells (SC), progenitor B cells (PROB), progenitor T cells (PRET), B cells (B), CD4 T cells (T4), CD8 T cells (T8), gamma-delta T cells (ɣδT), natural killer T cells (NKT), natural killer (NK), dendritic cells (DC), monocytes (MO), macrophages (MF), and granulocytes (GN).

We observed that neuro-receptors co-expressed in myeloid lineages were enriched for migratory functions and cell component movement including lysophosphatidic acid receptor (Lpar1) and purinergic receptors (P2rx4/P2ry12/P2ry6) known to be involved in myeloid cells migration^31,32^. Specifically enriched in DCs are neuropeptide Y receptor (Npy1r) known to promote DC trans endothelial migration^33^ and opioid receptor (Oprd1) which mediate chemotaxis in bone marrow-derived dendritic cells^34^. Endothelin receptor type B (Ednrb) and type A (Ednra) expressed in MF and GN (respectively) were enriched for migration^35^ and neutrophil chemotaxis^36^. Beyond functional enrichment, we noted other myeloid co-expressed neuro-receptors associated with migration functionality such as receptors of histamine^37^, glutamate^38^, serotonin^39^ and gastrin-releasing peptide^40^, strengthening the assumption that neuro-receptors play a role as migration regulators in myeloid cells. Neuro-receptors co-expressed in stem cells were enriched for function in metabolic processes which dictate the hematopoietic differentiation and maturation^41^ including growth hormone receptor (Ghr)^42^, serotonin receptor 5-HT (Htr2a)^43^ and luteinizing hormone /choriogonadotropin receptor (Lhcgr)^44^. Other neuro-receptors co-expressed in progenitor cells: receptors for glutamate, acetylcholine and melanocortin are known to be involved in synaptic, neural and erythroblast differentiation^45–47^, and as such, may also regulate immune cells differentiation. In the adaptive system we identified neuro-receptors as negative regulators of the inflammatory response in B cells including β_2_-adrenergic receptor (Adrb2)^48^ and cannabinoid receptor (Cnr2)^49^. With an opposite effect, neuro-receptors were identified as positive regulators of T cells activation, proliferation, migration, and cytokine production including Sphingosine-1-phosphate receptor (S1pr1)^50^ and proteinase activated receptors (F2r, F2rl1)^51,52^. Neural signaling in lymphocyte activation and inhibition provide a mechanism to “fine-tune” the adaptive response. Thus, this landscape of co-expressed neuro-receptors with lineage specific expression, creates a coordinated network of genes functioning in common processes of the immune system.

### Neuro-receptor expression profile recapitulates hematopoietic cellular differentiation

Immune cells originate from hematopoietic stem cells (HSCs) through a developmentally regulated process that gives rise to all immune cell lineages. The fate of HSC is modulated by the types and levels of signals used to stimulate them and by receptors’ expression dynamics^53,54^. To understand the influence of neural signals on immune lineage commitment and differentiation we generated an unsupervised trajectory of the stem cells and progenitor cells in the bone marrow and thymus based on the expression of neuro-receptors (see Methods). This data-driven analysis yielded a developmental trajectory which begins in stem cells and branches to two different paths of T and B progenitor cells [Figure 6A]. Repeating the trajectory building process for T cells and B cells separately we observed that the cells are ordered by the exact stages of differentiation [Figure 6B for B-cells, Figure S6 for T-cells]. Taken together these results suggest that neuro-receptor expression profile can recapitulate the hematopoietic cellular differentiation and highlight their potential functional importance in this process.

**Figure 6:**
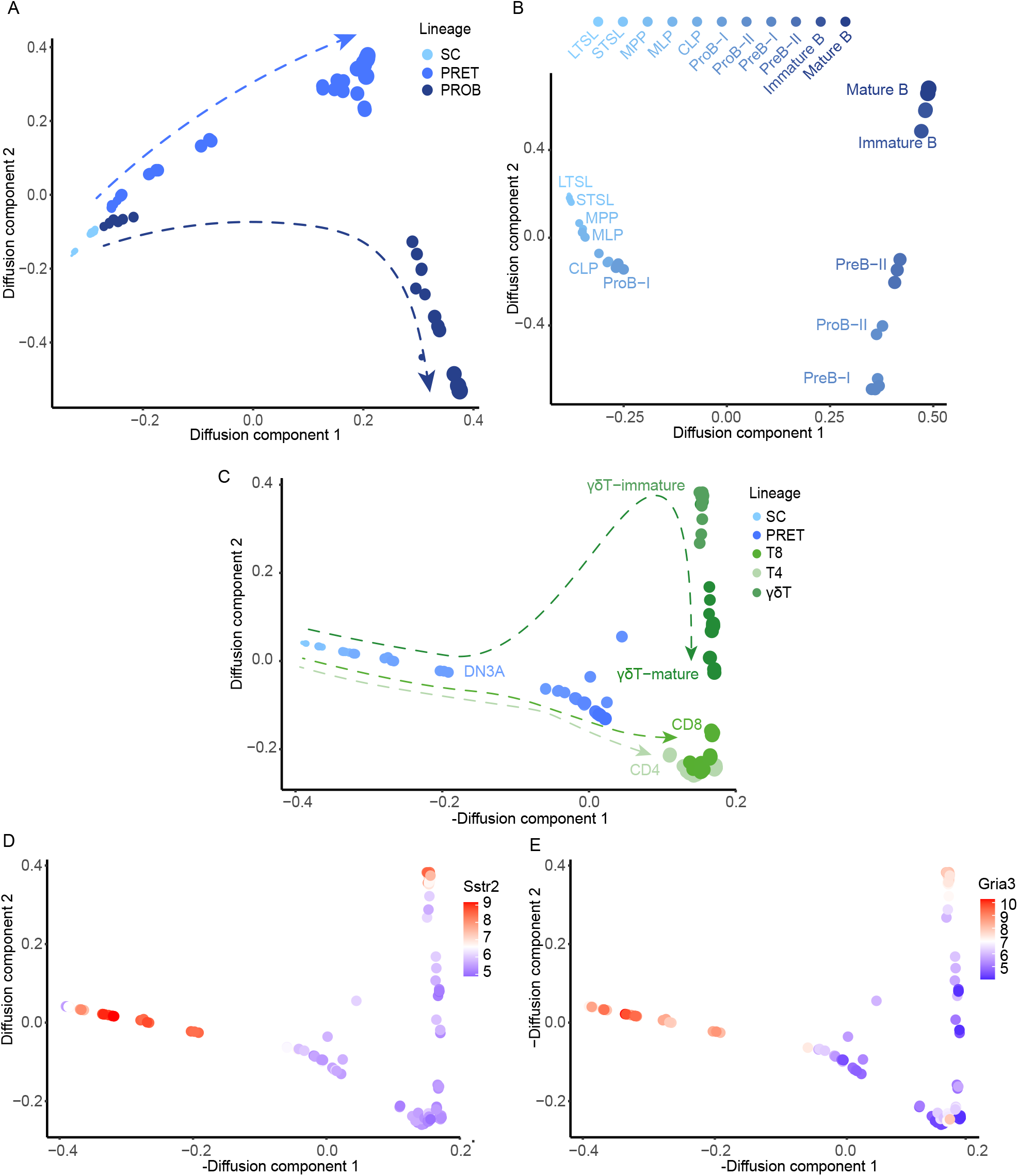
Neuro-receptor expression profile recapitulates hematopoietic cellular differentiation. Unsupervised developmental trajectories was generated using diffusion maps algorithm applied on immune progenitor cells at different stages of differentiation based on log2 expression profile of neuro-receptors differentially expressed (one-way ANOVA with *p-value <0*.*01*) between the progenitor cell types analyzed (in replicates). **(A)** Application of diffusion maps to 64 neuro-receptors differentially expressed between 25 cell subsets during early hematopoiesis stages sorted from bone marrow and thymus. Color indicates immune cell lineage. The arrows illustrate the differentiation trajectories direction which begins in stem cells and branches to two different paths of T and B progenitor cells. **(B)** Diffusion map applied on gene expression of 48 neuro-receptors differentially expressed between 11 progenitors B cell subsets from bone marrow, creating a trajectory of B cells ordered by the exact stages of differentiation (top) described by gradient color. **(C)** Diffusion map applied on gene expression of 61 neuro-receptors differentially expressed between 40 progenitor and mature T cell types from bone marrow and thymus. Color indicates immune lineage with gradient color and size representing the progenitor cells differentiation stages. The arrows illustrate the differentiation trajectories direction showing merged trajectories for T4 and T8 cells and branched trajectory of ɣδ T cells at the stage of double negative 3A (DN3A). (**D, E**) Diffusion map of T cells described in panel C, cells are colored by the log2 expression of Gria3 and Sstr2 respectively, showing their influence on immature ɣδ T cells commitment and their downregulation in mature ɣδ T cells.

The thymus maintains an environment critical for promoting T cells development and selection of the T cell repertoire. To test the role of neural signals in this process we added to the trajectory generation the developmental stages of CD4 and CD8 T cells and ɣδ T cells in the Thymus [Figure 6C]. In the T cells trajectory, the developmental stages of CD4 and CD8 T cells merge indicating that their commitment is not regulated by neural signaling. In contrast, the commitment of ɣδ T cells was captured in the neuro-receptor driven trajectory as immature ɣδ T cells diverged from the main trajectory branch at DN3a stage, influenced by the expression of somatostatin receptor (Sstr2) and glutamate receptor (Gria3) [Figures 6D,E]. These neuro-receptors are expressed in stem cells throughout all stages of immature ɣδ T cells and are downregulated in mature ɣδ T cells, suggesting their effect on ɣδ lineage commitment and maturation. Of note, we observed a gap between DN3a and DN3b in the thymus [Figure 6C, Figure S6], pointing to a possible regulatory transition by neural signals at the stage of beta selection checkpoint, associated with the major alterations in gene expression and increased proliferation occurring at this stage^55^. Thus, neuro-receptors expression changes allow ordering of HSC differentiation stages and commitment to specific lineage and lineage subtypes, suggesting that neural signals serve as differentiation regulators at critical stages of development.

### Neuro-receptors function is synchronized to regulate human B cells development

To integrate our findings, we focused on the B cells differentiation dynamics, where we hypothesized neuro-receptors function together along the developmental trajectory, playing a major part in the construction of immune-competent repertoire of naïve B-cells. Clustering neuro-receptor expression profiles in progenitor B cells yielded four modules describing co-regulated genes with similar behavior along B-cell development [Figure 7A, Figure S7A,B]. Neuro-receptors co-expressed at early stages of hematopoiesis possibly take part in B cell precursors commitment (modules 1,2 and 3) and neuro-receptors co-expressed at the later stages of differentiation might take part in the proliferation and maturation of B cells (module 3,4). In modules 2 and 3 neuro-receptors expression peak at stage Pro-B-I, where BCR heavy chain immunoglobulin genes rearrangement begins. Here too we observed a gap in B cells trajectory after this stage [Figure 6B] suggesting a regulatory transition where the neural signals monitor for functional Ig H-chain rearrangement and trigger clonal expansion and developmental progression^56^. To validate our findings, we analyzed the neuro-receptors expression at the protein level and tested their clinical translation to human. We performed a mass cytometry (CYTOF) single cell-proteomics analysis on human bone marrow samples from three healthy donors, analyzing a subset of neuro-receptors from our findings. We observed broad agreement with the dynamics we inferred from our analysis of mouse [Figure 7B, Figure S7B]: Proteinase activated receptor F2R (Module 1), shown to regulate the HCS retention in bone marrow^57^, was indeed expressed at early stages of progenitor B cells (adjusted *P* =1.8E-5). Glutamate receptor GRIA3 (Module 2) dynamics peaked at stage Pro-B-I and then decreased sharply (adjusted *P*=0.003), in agreement with our mouse analysis. Though GRIA3 function in B cell development has not been ascribed, its family member GRIA1 is associated with human megakaryocyte maturation^58^. Examining neuro-receptors that were upregulated along the mouse B cells trajectory, β_2_-adrenergic receptor ADRB2, and cannabinoid receptor CNR2 (both in module 4) were indeed upregulated at the later stages of differentiation in human trajectory (adjusted *P* = 0.09, 0.3 respectively). Of relevance, β_2_-adrenergic receptors have been shown to play a role in B cell progenitor egress from the bone marrow ^59,60^ and CNR2 is required for the retention of immature B cells in the sinusoids upon egress from the bone marrow parenchyma^49^. Arginine vasopressin receptor AVPR2 (in module 4) did not show reproducible results in human bone marrow samples and the expression did not increase gradually as we observed in mouse mRNA. Our experimental results highlight the relevance of neural signals as regulators of the hematopoietic cellular differentiation process and in the context of our system level mapping the importance of considering neural signals as key players in immunology.

**Figure 7:**
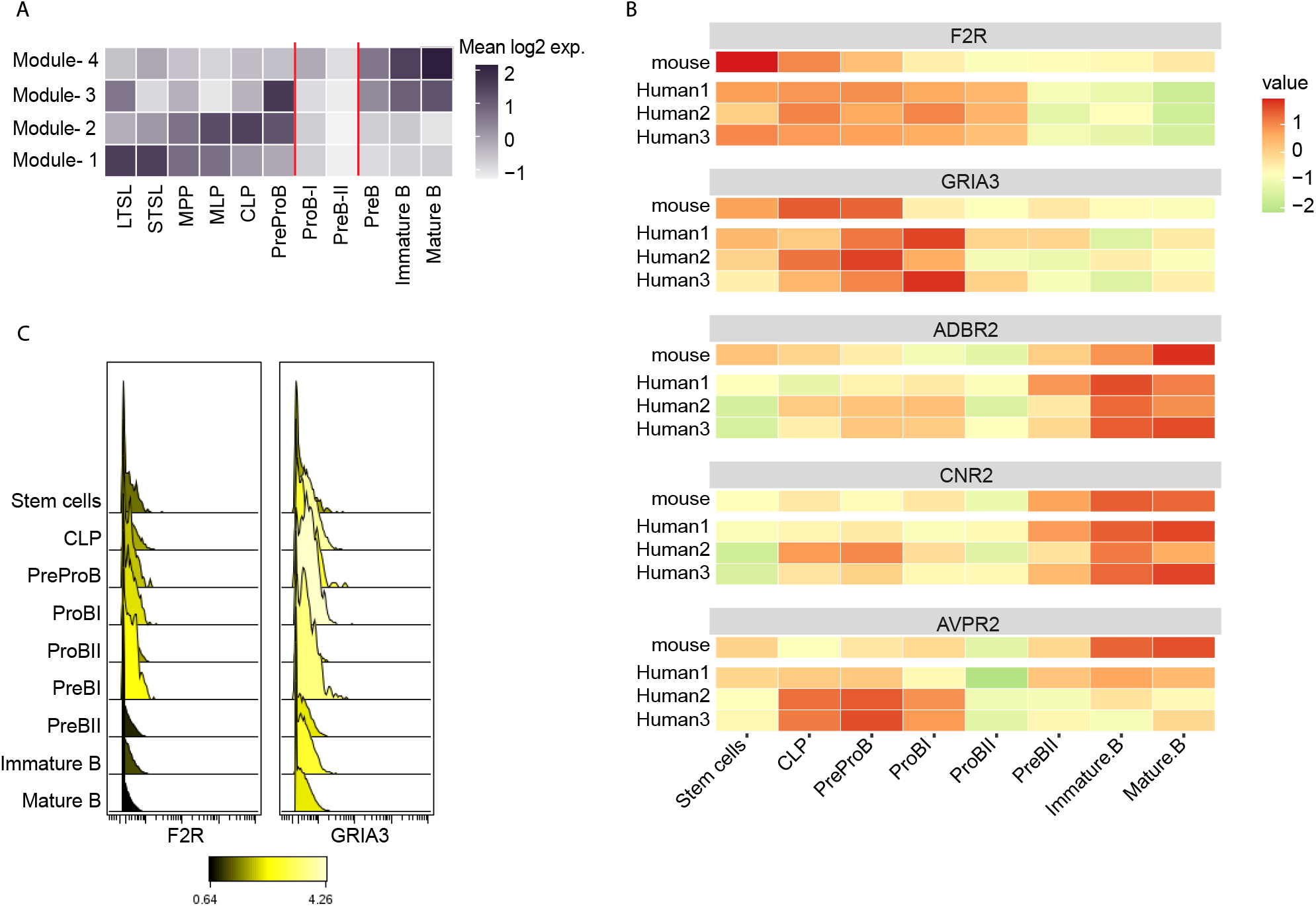
Neuro-receptors function together to regulate human B cells development. Neuro-receptors are co-expressed along different stages of the developmental trajectory. (**A**) Heatmap of the mean log2 expression of neuro-receptors clustered to 4 modules along the B cells trajectory by WGCNA. The cells are ordered by the differentiation stages. Red lines highlight major stages of change in modules expression. (**B, C**) Neuro-receptor protein expression in human bone marrow b-cells follows hematopoiesis and validates mouse mRNA prediction. (**B**) Heatmap indicating the protein expression of each neuro-receptor analyzed by CyTOF analysis in 3 human bone marrow samples along B cell differentiation populations ordered by their differentiation stages. The mouse equivalent B-cell mRNA for the gene is shown for comparison. Proteins analyzed are: Proteinase activated receptor (F2R), Glutamate iontropic receptor (GRIA3), β2-adrenergic receptor (ADRB2), cannabinoid receptor (CNR2) and arginine vasopressin receptor (AVPR2). Each protein expression is transformed to the 90th percentile of the marker expression in each annotated cell population, normalized by the protein minimum value and scaled. (**C**) Example of neuro-receptors F2R and Gria3 extracellular staining intensity (CyTOF) on progenitor B cell types defined with a histogram. Color indicates protein median intensity in the cell population.

## Discussion

Accumulating evidence in recent years indicate a strong link between neural signals and the immune system. This observation invoked us to systematically explore the neuro-immune regulation network. Our gene expression analysis identified 122 neuro-receptors and 38 neural ligands (51% and 67% respectively) implicated in the peripheral immune system in steady state. This is a much broader milieu than appreciated to date that is in line in terms of communication complexity to that of cytokines and chemokines. We show that studying individual canonical neural signals as a model to describe inflammatory regulation is restrictive, whereas studying the combined effect of the full set of neural signals highlights their importance in immunology.

A global map of neuro-immune communication revealed a specific regulatory structure where the transcriptional profile of neuro-receptors across immune cell types is highly specific to lineage. We identified that pDCs and ɣδ T cells are highly specialized in their expressed set of neuro-receptors, possibly because the interface they play between innate and adaptive immunity makes them important targets of neuronal regulation, orchestrating the entire immune system. Further analysis of single-cell RNA-seq data may identify additional neuro-receptors expression in rare cell types though these may be limited in insight due to low abundance expression detection being a known issue in this platform^24^.

Analyzing neuro-receptor co-expression, we observed a coordinated network of genes working together with cytokines receptors and functioning in shared processes such as migration, chemotaxis, development, proliferation, and activation of inflammatory processes. We observed a specific migratory effect of neural signals on immune cells in the innate immunity branch-the first line of defense, where the speed of response and migration is important for successfully protecting the host. During pathogen detection, neurons are activated faster than immune cells^24^, and so the rapid release of neural mediators and the expression of neuro-receptors on immune cells can act to accelerate immune response for host protection. Furthermore, neuro-receptors with unknown function in the immune system were associated with biological processes enriched within their cluster, such as migration and differentiation, expanding knowledge of the regulatory network of neural signals in the immune system. Further characterization of neuro-receptors pro-inflammatory or anti-inflammatory role will create a clearer picture of the neural signaling role in maintaining a balanced inflammatory response in homeostasis.

The major source of neural signaling molecules are neurons; however, many “neural signals” are produced also by non-neural cells including immune cells, taking part in immune intrinsic communication, and functioning as bi-directional communication mediators between the different systems. Expanding our search to the immune microenvironment, we identified a high percent of neural genes shared between immune and stromal cells which are key regulators of the immune system. This emphasizes the role neural signals play as a common communication language between the nervous system, the immune system and the microenvironment in a tissue dependent manner, especially with respect to regulation of resident immune cells. Reports in the literature indicate that immune regulation is dependent also on the location in the tissue^61^, which we observed for the different immune populations in the gut having protective vs. immunosuppressive functions. This phenomenon should be further investigated using spatial ‘omic and imaging approaches. Similarly, the tri-directional immune-neuronal-stromal communication network can be extended to cover immune activation states and specifically neuro-inflammatory disorders in the periophery.

The discovery that neuro-receptors are broadly expressed in immune cell types in a non-uniform distribution and with specific functions, strongly points to a high selection pressure for cell-specific neural regulation. Beyond their role as lineage specific regulators, the immune cells sensitivity to neural signal transduction within lineage is modulated through differential expression of the neuro-receptor subtypes. We detected selective functionality for many neuro-receptors in the complex life-cycle of leukocytes, with a clear effect on the differentiation and proliferation of the immune cells. An exciting observation stemming from our data is that the neuro-receptors expressed on hematopoietic stem cells and progenitor cells recapitulated the hematopoiesis developmental trajectory. By a system level view of neuro-receptors along the B cells differentiation process we identified that the regulation by neural signals is achieved in two phases: a first set of neural signals regulating the early development phases of commitment and immunoglobulin rearrangement and a second set of neural signals that regulate the latter stages of survival, proliferation, and maturation. We also identified association between neuro-receptors expressed in stem cells to metabolic pathways known to regulate HSC function and fate decisions^41^, suggesting neural signals may affect metabolic adaptation of HSCs in differentiation. By these means, the neural signals probably regulate and navigate the host energy resource, maintaining most HSC in a quiescent state within their niche in steady state, and in response to inflammatory stressors elicit an increase of the entire repertoire of hematopoietic cells. These findings indicate neuro-receptors play a yet unappreciated role in hematopoiesis and may offer new directions for successful recapitulation in-vitro. Beyond hematopoiesis, the high selective pressure on cell specific expression and function suggests that the neurogenic and immune mediated inflammation are not independent entities but rather a joint system that evolved to protect the body and maintain homeostasis.

We provide here a resource of the potential neural molecular pathways supporting the regulation of the immune system at steady state. We envision this map of neuro-immune interactions will allow to approach neural signals as an integral part of the immune regulatory system and provide clear testable new avenues of experimental follow up for neuroimmunologists and immunologists alike. Broadly, our results reveal a broader alphabet than appreciated to date, which enables a richer vocabulary for immune social architecture^78^. Given the origin of these signals may come from the central nervous system, the system-level map we present here may serve to decipher how the brain monitor and controls immune response to emotions, stress and other perceptions.

## Acknowledgements

We thank members of the Shen-Orr lab for illuminating discussions and feedback. A. Rolls for guidance, A. Alpert and R. Normand for assistance with gene specific threshold development. D. Cohen for computing help, A. Grau for support with the CYTOF machine. Y. Avraham and Y. Schumacher for figure design and M. Lukacisin and R. Bendayan for manuscript comments.

## Author contributions

K.R.B. and S.S.S.-O. designed the overall data analysis strategy. K.R.B performed the overall data analysis, analyzed the cell subset data and gene expression data. N.M. assisted with the biological interpretation. N.M. M.S and E.S. performed the experimental validation. K.R.B, N.M. and S.S.S.-O. co-wrote the paper.

## Methods

### ImmGen data

We focused on immune cells isolated from mouse in steady-state conditions. The data includes 180 cell types across the entire murine hematopoietic lineage at major stages of differentiation as well as stromal cell, measured between 2-5 replicates generating a total of 567 samples. Cells are divided by annotation into 13 major immune lineages, including granulocytes (GN), monocytes (MO), macrophages (MF), dendritic cells (DC), natural killer (NK), gamma-delta T cells (ɣδT), natural killer T cells (NKT), CD8 (T8), CD4 (T4), B cells (B) and progenitor B cells (ProB), progenitor T cells (PreT) and stem cells (SC), as well as stromal cells [Table S1]. The samples were isolated from 20 lymphoid organs and other tissues in the periphery [Figure S1 for summary of the lineages isolated in each tissue]. All samples were hybridized with Affymetrix Gene ST 1.0 array platform, generated in 37 independent batches and normalized based on control samples as reported in detail by the ImmGen consortium^10^,^12^.

### List of neuro-receptors and ligands

To study the neural signals effect on the immune system we analyzed the expression levels of the receptors and ligands of neural signals in immune cells’ transcriptome. Member genes were identified based on a list of genes taken from KEGG pathway labeled ‘Neuroactive ligand-receptor interaction - Mus musculus’. These we narrowed down to a list consisting of 64 neural signals including 15 neurotransmitters, 31 neuropeptides and 18 neuro-hormones which we identified as well-known and specific neural regulators. Many neural ligands can bind to multiple receptors creating a grand list of 239 neuro-receptors [Table S2].

Neuropeptides and neuro-hormones are proteins, and their signal may be detected by gene expression data. Neurotransmitters, on the other hand, are chemical metabolites and their levels cannot be detected using gene expression. Thus, as an alternative solution, we estimated neurotransmitters activity by averaging for each neurotransmitter on every cell the expression of their precursors or enzymes involved in their final synthesis [Table S3].

To set a frame of reference we sought to assess the behavior of neural signals that have already been studied in the context of immune regulation. Specifically, we defined a set of canonical signals with known roles in the peripheral immune system including acetylcholine (ACh), norepinephrine (NE), dopamine (DA), neuromedin U (NmU), vasoactive intestinal peptide (VIP) and substance P (SP) ^3^.

### Expression threshold

ImmGen microarray data was log2 transformed and the expression value distribution range from approximately 3 to 12. Determining whether a gene is actually expressed, or not, in a given cell type requires setting a clear boundary between true expressions versus background signals. Towards this, we assessed two models:

### 1. Gaussian mixture model

The original ImmGen publication^12^ used an empirical Gaussian mixture model to evaluate probabilistic thresholds of expression. The log scaled expression histograms was decomposed into two Gaussian distributions using R package ‘mixtools’ [Figure S3A]. Based on the low distribution that corresponds to background noise, a threshold of 6.9 was set with 95% probability of true expression. Based on the higher distribution that corresponds to true signal, a threshold of 4.6 was set with 95% probability of the gene not being expressed. Of note, these two thresholds result in an intermediate range of expression values in which it is unclear whether the gene is expressed or not.

### 2. Gene specific threshold algorithm

A fixed global expression threshold across all genes, is a common practice. However, it is quite clear a threshold value appropriate for highly expressed genes may not be suitable for lowly expressed genes ^62^. We developed a model that determines for each gene individually, the best expression threshold that a gene is truly expressed at the range of intermediate probabilities defined by the Gaussian mixture model. The range of thresholds is restricted by a lower bound expression 4.6, as defined by pseudogenes, and an upper bound of 6.9 defining true expression, with bins of 0.1 resulting in a range of 24 threshold levels. The model is based on the observation seen in a subset of genes, for which the expression signal is enriched for a lineage population specific pattern [Figure 2A,B]. Broadly we define a gene specific threshold is based on the expression threshold for which a group of cell types sharing a common lineage are most enriched in the data [Figure S3C].

### Step 1- Detect gene enriched lineages

To test if the lineage cells are enriched compared to all ImmGen samples we applied a hypergeometric test. Specifically, we tested 24 expression threshold levels in the range of 4.6-6.9 log2 expression, converting the gene expression values for these set of thresholds to a binary vector representing a value of expressed (1) or not expressed (0). At each threshold the hypergeometric test [equation 1] measures the probability of having k cell types in the lineage expressing the gene (out of n cell types expressing the gene in the data) from N cell types in the data containing K cell types of the lineage.

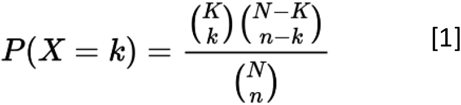

Lineage groups with adjusted p-values < 0.01 (using Benjamini & Hochberg (BH) correction) were considered significantly enriched.

Enriched lineages may cover all lineage cell types or a subset of cell types which are differentially expressed within the lineage. We defined lineage subset for lineages significantly differentially expressed (one-way ANOVA, Pr(>F) <0.01), by computing Tukeys honest significant differences (HSD) between each pair of cell types in the lineage. Cell types enriched in comparisons with adjusted pvalue <0.01 were included in lineage subset group. We tested the possibility that lineage subsets are enriched compared to ImmGen data by applying a hypergeometric test at every value in the threshold range, followed by a Benjamini & Hochberg correction. We considered the lineage group of cell types (lineage subset or all lineage cell types) meeting the minimum hypergeometric adjusted p-value at any threshold as the lineage enriched group. Lineages with no significant enrichment at any threshold were not included in the next step.

### Step 2- Select optimal threshold with the most significant enrichment

We combined all groups of lineages found enriched in Step 1 and computed hypergeometric tests at 24 expression thresholds. We defined the gene specific threshold to be at the expression level where the combined group is over expressed compared to all ImmGen samples with the minimum hypergeometric p-value.

### Step 3- Examining reliability of gene specific threshold

The gene-specific threshold procedure we described above allows to ‘rescue’ cell types which would have been declared as not expressing a gene based on the global expression threshold of 6.9. We performed a cleanup step to ensure that at gene specific threshold the cell types rescued are from the enriched group and not randomly added samples. We calculated the ratio between cell types from enriched group that were rescued compared to all other cell types rescued. Genes with ratio value smaller than 0.5 were considered unreliable with FDR > 8.9% [Figure S3B] and the gene-specific threshold was replaced with the general threshold of 6.9.

### Expression statistics

Genes with expression values above their specific gene threshold were defined expressed [Table S2]. Cell types found expressed in the intermediate range would not have been expressed using the fixed threshold of 6.9 and there for are rescued using the gene specific threshold approach [Figure 2C]. The specificity of the genes is described by the number of lineages it is expressed in [Figure 2D].

### Principle component analysis

We visualized and assessed similarities and differences between cell types based on the expression profile of all neuro-receptors. PCA was computed by the R function prcomp with scale parameter set to false. We examined whether the immune cells are grouped together by specific phenotype, evaluating the relationships at the lineage and tissue level. We performed PCA based on expression profile of: 1-all neuro-receptors, 2-canonical neural signals receptors, 3-cytokine receptors.

Separation measure: To compare the multivariate distance between the distribution of progenitor cells, lymphoid and myeloid cells in the first two PCs we used Mahalanobis distance which tests the mean difference between every 2 cell populations while considering the covariance. The significance of separation (pValue) was calculated by Hotelling T2 statistics. The standard error for each comparison was estimated based on bootstrapping the data 100 times, sampling with replacement the cells in PCA and then calculating Mahalanobis distance.

Controlling for gene set size: We verified that the differences in immune group separation in PCA observed between neuro-receptors and canonical neuro-receptors or cytokine receptors is not due to the difference in gene set size. We sampled neuro-receptors with gene set sizes of canonical neuro-receptors list and cytokine receptors list (41 and 110 respectively) 100 times each, and calculated the Mahalanobis distance. We calculated the significance in differences in Mahalanobis distance between the different analysis (neuro-receptors, canonical neuro-receptors and cytokine-receptors) using t.test.

PCA based on random gene set: We evaluated immune cell groups separation in PCA based on neuro-receptors compared to random sets of genes. We sampled 240 genes randomly from the transcriptome discarding neuro-receptors and cytokine receptors. We calculated Mahalanobis distance between immune cell groups of lymphocytes, myeloid and progenitor cells, sampling the data 100 times. We calculated the significance in Mahalanobis distance differences between neuro-receptors and random gene sets using t.test.

PCA of dendritic cells (DCs): We analyzed the landscape of DCs with PCA described by 122 expressed neuro-receptors. In order to identify the neuro-receptors that mostly contribute to the variability and to the observed separation of plasmacytoid DCs (pDCs) in the second principle component (PC2) we selected neuro-receptors with weights < -0.2 in the linear combination of PC2 [Figure 4D].

### Tissue communication network assembly

To interpret cell to cell communication via neural signals we constructed networks of ligand-receptor interactions between the different cell types in each tissue separately. We obtained ligand-receptor pairs from KEGG pathway ‘Neuroactive ligand-receptor interaction - Mus musculus’. We assembled the interaction networks based on the binary expression matrix of neuro-receptors and ligands in immune and stromal cells in ImmGen data (based on the gene specific threshold).

### Intestinal neuro-immune cell level communication network

In the network of neuro-immune interactions in the intestine we integrated sorted and single-cell data to obtain interactions of immune cells expressing neuro-receptor to enteric neurons expressing the corresponding neural ligand. We analyzed enteric neurons from single cell data set classified as cholinergic, glutaminercis and nitrergic^15^. To associate neuronal cell types with expressed neural ligands we used the data set Trinarization scores defining if gene is expressed or not^15^. We extended the intestine network to include neural communication pathways within the immune system, involved in immune autocrine or paracrine regulation. We used CytoScape to visualize the network ^63^.

### Cell’s specialization and Diversity of neuro-receptors expression profile

We characterized the immune cells diversity and specialization of neuro-receptors expression profile using functions based on Shannon’s information theory26 [Figure 4A]. The cell diversity Hj was computed by the R function ‘entropyDiversity’ which is an adaptation of Shannon’s entropy formula to the transcriptome frequencies:

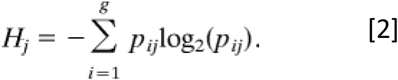

were *pij* is the relative abundance for the *i*th gene in the *j*th cell type. The highest diversity score is given when the relative abundance of all genes is 0.5, meaning cells with higher diversity correspond to a more uniform distribution of neuro-receptor expression frequencies. In Microarray platform the expression data is not absolute, and the genes may have a uniform distribution at expression levels above expression threshold or at the background levels of expression below threshold. This means high diversity may correspond to one of two extreme states: either cells expressing a large fraction of neuro-receptors with expression levels above threshold, or those cells expressing most neuro-receptors below expression threshold.

Cell specialization *δj* [equation 4] was quantified using the R function ‘sample Specialization’ that measures the average gene specificity *Si* which is a linear function of the entropy of a gene’s expression distribution [equation 3].

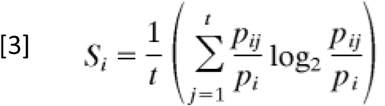

were *pi* is the average expression of the *i*th gene.

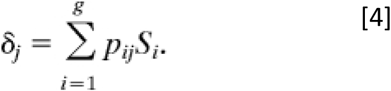

From equations 3 and 4 we can see that the specificity and specialization are sensitive to the number of samples expressing the gene, and the specificity of neuro-receptors enriched in a lineage is dependent on the number of cell types in the lineage. Therefore, we excluded GN cells from this part of the analysis because of very few cell types (2). The specificity and diversity were calculated for 168 cell types based on the expression profile of 112 expressed neuro-receptors. The specialization and diversity scores were scaled between 0 - 1. We tested the association between the cells diversity and specialization by calculating Pearson correlation coefficient for all immune cells and separately for each lineage cell types. The centroid of each lineage was calculated as the lineage cell types mean diversity and specialization.

To evaluate if the negative correlation between immune cells diversity and specialization is a unique property of neuro-receptors expression we compared it to the correlation of immune cells diversity and specialization scores based on 105 expressed cytokine receptors expression profile.

### Diffusion map analysis

To test if the high-dimensional neuro-receptors expression relations between hematopoietic progenitor cells construct a continuous transition of the cells by the developmental stages (developmental trajectory), we performed dimensionality reduction using the diffusion map method (R package destiny ^64^) with k=20. To reduce noise, we only included the neuro-receptors expressed (above their gene specific threshold) and with a significantly different expression pattern between the subset of progenitor cells analyzed (one way ANOVA with p-value <0.05).

### Co-expression analysis

Expressed neuro-receptors were clustered to modules of coordinated expression pattern across immune cell types using unsupervised weighted gene co-expression network analysis (WGCNA) R package ^63^. We construct a signed network by calculating Pearson correlation coefficients for all pairs of neuro-receptors and used the scale-free topology criterion with a choice of ß =10.

All immune cells: WGCNA assigned 122 expressed neuro-receptors to 9 modules with similar expression profiles containing 3 to 74 genes using average linkage hierarchical clustering [Table S5].

B cells differentiation: WGCNA on 11 cell types of stem cells and progenitor B cells in the bone marrow assigning 48 neuro-receptors to 4 modules with similar expression profile along the B cells differentiation process [Figure 7A, Figure S7A,B].

Module-lineage relationship: For each module we calculated the Module Eigengene (ME), defined as the first principal component of the module which represents the expression profile of the module. We evaluated the correlation between MEs and the cells lineage trait represented as binary vector, using Pearson’s correlation coefficients and correlation test p-value [Figure 5A].

Receptor – lineage relationship: Pair-wise correlation between receptor expression profile (neuro-receptors and cytokine receptors) and the sample lineage binary vector was conducted to identify genes with significant correlation to lineage trait (Bonferroni correction < 0.05). A visualization of the 122 neuro-receptors and 105 cytokine receptors significantly correlated to lineage was conducted using ‘circlize’ R package^65^ [Figure 5B].

Functional enrichment analysis: To evaluate if co-expressed genes enriched to lineage are also active in the same biological processes we tested for over-representation of Gene Ontology (GO) terms using clusterProfiler package^66^. enrichGO function was run on two unranked lists of genes where the background list was given as all the expressed neur-receptors in our analysis [Figure 5C, Tables S6,S8].

### CyTOF analysis

Frozen human bone marrow mononuclear cells (BMMC) from three healthy donors were purchased from STEMCELL Technologies and stored in liquid nitrogen until use. Upon thawing, cells were immediately washed with warm RPMI containing 20% FCS, Pen Strep Glutamine and DNaseI (Pierce™ Universal Nuclease for Cell Lysis, Thermo Fisher Scientific) to prevent cell clumping. Next, 1.2×10^7^ cells were incubated with Cell-ID™ Intercalator-103Rh (Fluidigm Inc.), for live-dead cell discrimination as per manufacturer’s instructions. Cells were then washed twice by centrifugation (600g, 5 min) with Maxpar® Cell Staining Buffer (CSM, Fluidigm Inc.) and stained with a panel of metal-tagged antibodies designed to detect in high resolution B lymphocytes bone marrow differentiation stages in addition to antibodies against neuro-receptors implicated in B cell differentiation (complete list of antibodies and their catalog numbers is provided in Supplementary Table 8). Antibodies for immune-phenotyping were validated by the manufacturers for Mass cytometry application (as indicated on the manufacturer’s datasheet, available online) and were metal conjugated using the MAXPAR reagent (Fluidigm Inc.). BMMC were first stained for surface epitopes with an extracellular mix (Supplementary Table 8) for 1 h at room temperature. Cells were washed by centrifugation (600g, 5 min) with CSM, and then fixed in 1.6% paraformaldehyde (Thermo Scientific™ Pierce™). Cells were then permeabilized by cold methanol and stained with an intracellular mix for 30 min at room temperature followed by a wash and incubated in 1.6% paraformaldehyde at 4°C overnight. The following day, cells were stained Cell-ID™ iridium Intercalator-to identify single cells, washed, resuspended with Maxpar Cell Acquisition Solution (Fluidigm, Inc.) and acquired by Helios Mass Cytometer (Fluidigm) at approximately 300 events per second. A minimum of 3×10^6^ cells were acquired for each donor to allow the detection of rare B lymphocyte differentiation subsets. Files were then normalized per donor using premessa R package and analyzed by CytoBank (Kotecha N, Mende). Cell populations were gated manually [Figure S8] and the neuro-receptors expression was defined based on the 90^th^ percentile in the corresponding immune population. The data was scaled using ‘Transformed ratio’ compared to column minimum expression.

### Neural genes expressed in immune vs. stromal cells

We compared the expressed neuro-receptors and ligands in any immune cell type across all 19 tissues to the expressed neuro-receptors and ligands in stromal cells in the tissues: mesenteric lymph node, thymus and skin. We then analyzed how many expressed genes are shared between immune cells and stromal cells in the peripheral tissues. We note this comparison is likely a lower bound it does not include all stromal cell types and all peripheral tissues.

