## Supplementary Figure for "Systematic mapping of neural signals in the periphery reveals a rich and specific alphabet for immune cell communication"

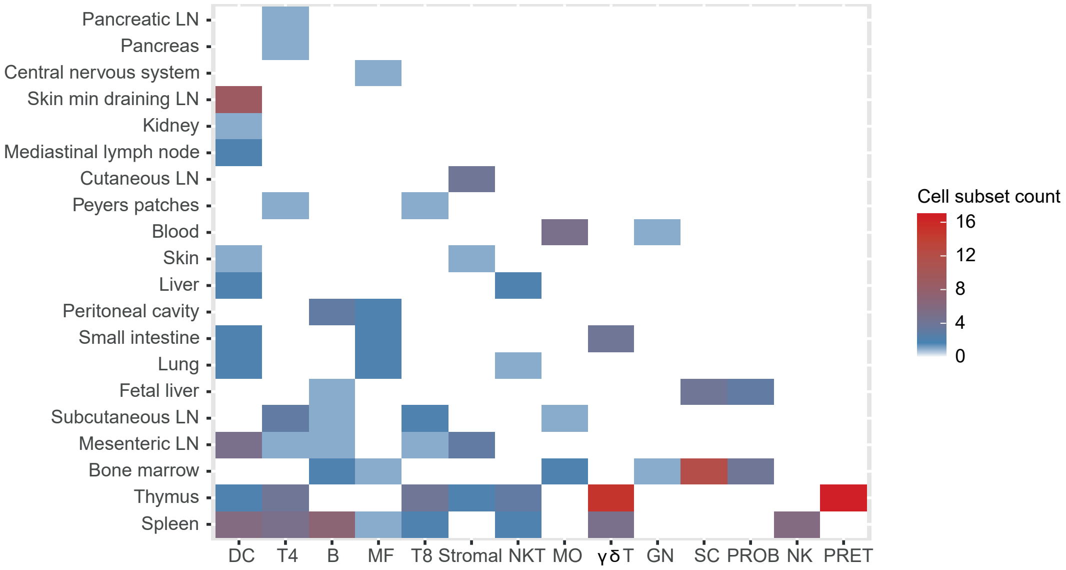


**Figure S1: Summary of the repertoire of immune samples in ImmGen dataset.** Heatmap indicating the number of immune cell types by lineage (x axis) and tissue of origin (y axis). Lineages and Tissues are ordered by the collective number of immune cell types they include. Of note, immune lineages are not isolated uniformly from all tissues. Lineages included: stem cells (SC), progenitor B cells (PROB), progenitor T cells (PRET), B cells (B), CD4 T cells (T4), CD8 T cells (T8), gamma-delta T cells (ɣδT), natural killer T cells (NKT), natural killer (NK), dendritic cells (DC), monocytes (MO), macrophages (MF), granulocytes (GN) and Stromal cells. LN- lymph node.

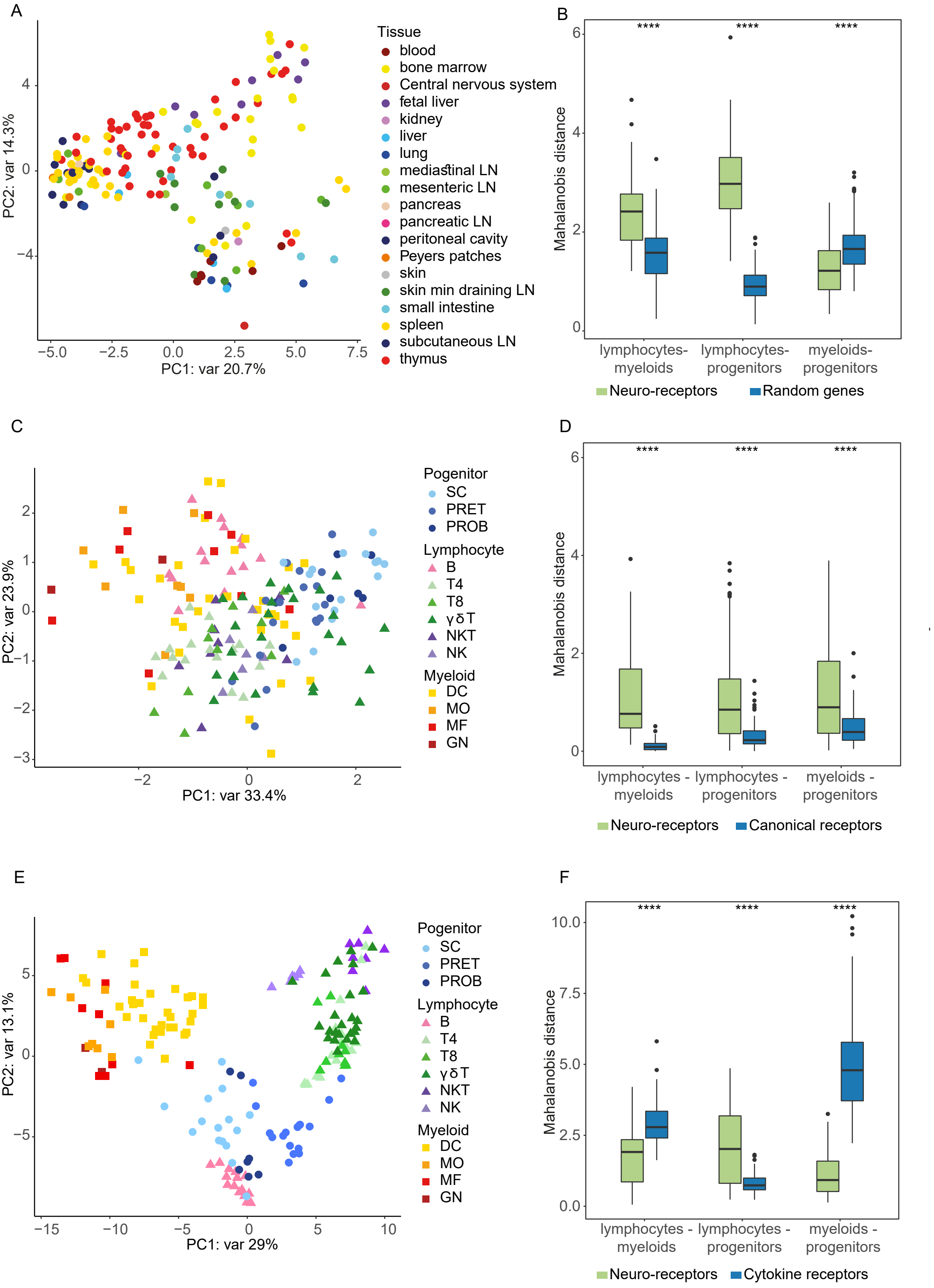
A
B
C
D
E
F
PC1: var 20.7%
PC2: var 14.3%
Tissue
blood
bone marrow
Central nervous system
fetal liver
kidney
liver
lung
mediastinal LN
mesenteric LN
pancreas
pancreatic LN
peritoneal cavity
Peyers patches
skin
skin min draining LN
small intestine
spleen
subcutaneous LN
thymus
Mahalanobis distance
****
lymphocytes-myeloids
lymphocytes-progenitors
myeloids-progenitors
Neuro-receptors
Random genes
PC1: var 33.4%
PC2: var 23.9%
Pogenitor
SC
PRET
PROB
Lymphocyte
B
T4
T8
γδT
NKT
NK
Myeloid
DC
MO
MF
GN
lymphocytes - myeloids
lymphocytes - progenitors
myeloids - progenitors
Canonical receptors
PC1: var 29%
PC2: var 13.1%
Cytokine receptors

**Figure S2: Neuro-receptor expression profile distinguishes between immune lineages. (A)** PCA based on neuro-receptors log2 expression for all immune cell types (mean of replicates). Color indicates the tissue of origin of sorted cells. **(B)** Bar plot comparing the multivariate distance in PCA space between lymphocytes, meyloids and progenitor cell subsets, analyzed based on either neuro-receptors (green) or random sets of genes sampled 100 times (blue). Mahalanobis distance standard error for neuro-receptors analysis was estimated based on bootstrapping the data 100 times. **(C)** PCA based on canonical neural signals' receptors log2 expression for all immune cell type (considering acetylcholine, norepinephrine, dopamine, neuromedin U, vasoactive intestinal peptide, substance P and calcitonin canonical neural signals). Color indicates cell lineage and shape indicates lineage type: lymphocyte, meyloid or progenitor. **(D)** Bar plot comparing the multivariate distance in PCA space between lymphocytes, meyloids and progenitor cell subsets analyzed based on either neuro-receptors (green) or canonical neuro-receptors (blue). We controlled for group size by subseting neuro-receptors to 41 genes 100 times. Mahalanobis distance standard error for canonical neuro-receptors analysis was estimated based on bootstrapping the data 100 times. **(E)** PCA based on 110 cytokines receptors mean log2 expression for all immune cell type. Color indicates cell lineage and shape indicates lineage type: lymphocyte, meyloid or progenitor. **(F)** Bar plot comparing the multivariate distance in PCA space between lymphocytes, meyloids and progenitor cell subsets analyzed based on either neuro-receptors (green) or cytokines receptors (blue). We controlled for group size by subseting neuro-receptors to 110 genes 100 times. Mahalanobis distance standard error for cytokines receptors based PCA was estimated based on bootstrapping the data 100 times.

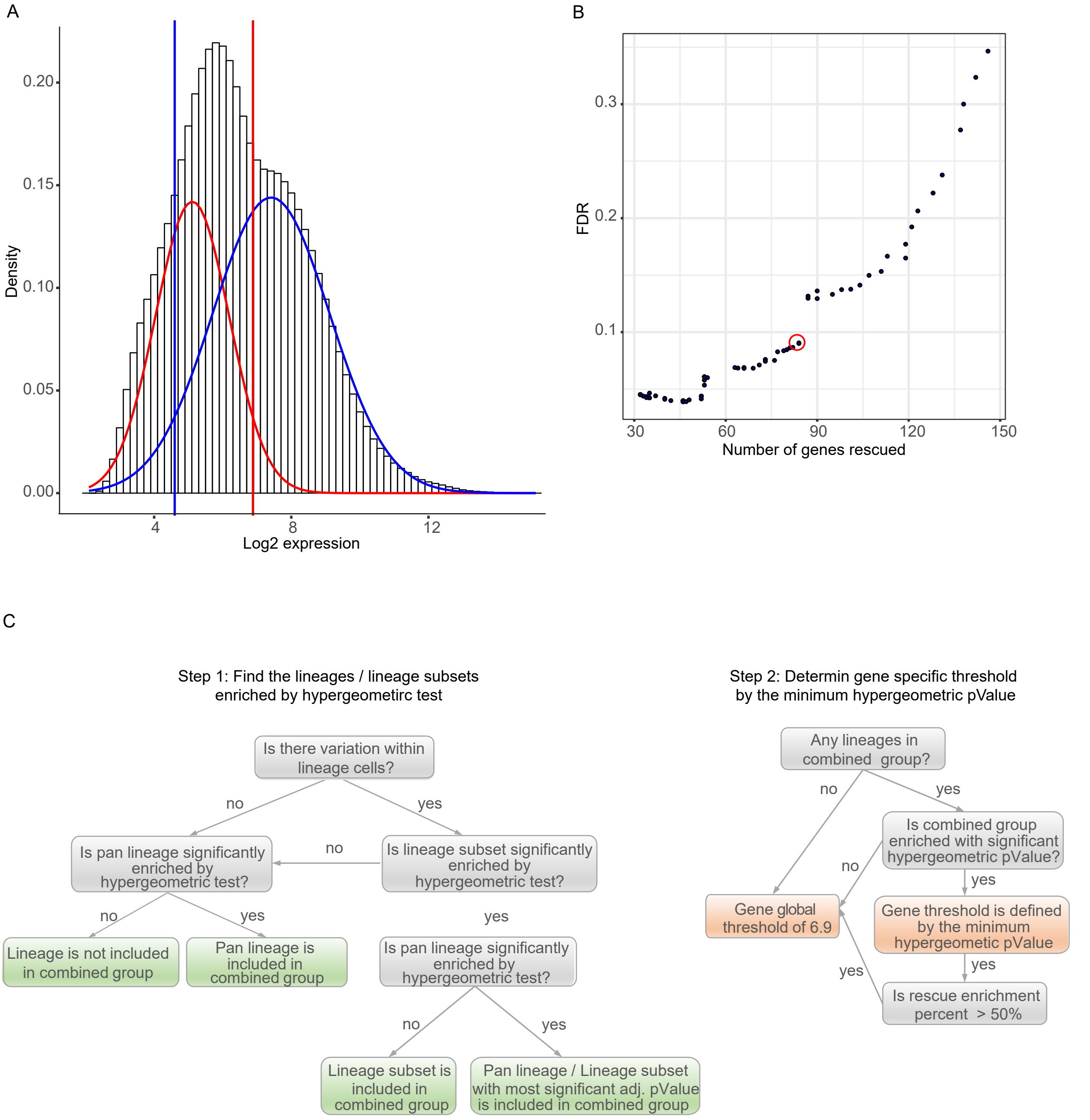


**Figure S3: A need for gene specific threshold for determining expression.** (**A**) Determining gene expression global threshold based on Gaussian mixture model used by the original ImmGen publication. The distribution of log2 expression values decomposed into the sum of two Gaussian distributions. The red line at log2 expression of 6.9 indicates the global threshold of expression computed from the lower (red) distribution, and the blue line at log2 expression of 4.6 indicates the threshold of background signal computed from the higher (blue) distribution. Between these two lines is the intermediate expression range for which it is difficult to determine with high probability if truly expressed or not. (**B**) Setting threshold for unreliable gene specific thresholds. Scatter plot of FDR vs. number of genes rescued. We calculated ratio score of cell types rescued from enriched group to all rescued cells, and defined FDR as the ratio score calculated for neural genes vs. mean ratio score calculated based on permutations. Marked in red circle is the threshold chosen. (**C**) Decision tree describing gene specific threshold algorithem: Step 1 was conducted for each lineage seperately, determining if the expression of the gene is enriched in the lineage and determine if the pan lineage or lineage subset will be included in the combined group in step 2 (green boxes). Steps 2 combine all the lineages cell goups found enriched in step1 as combined group, and identifies the best gene specific expression threshold (orange boxes).

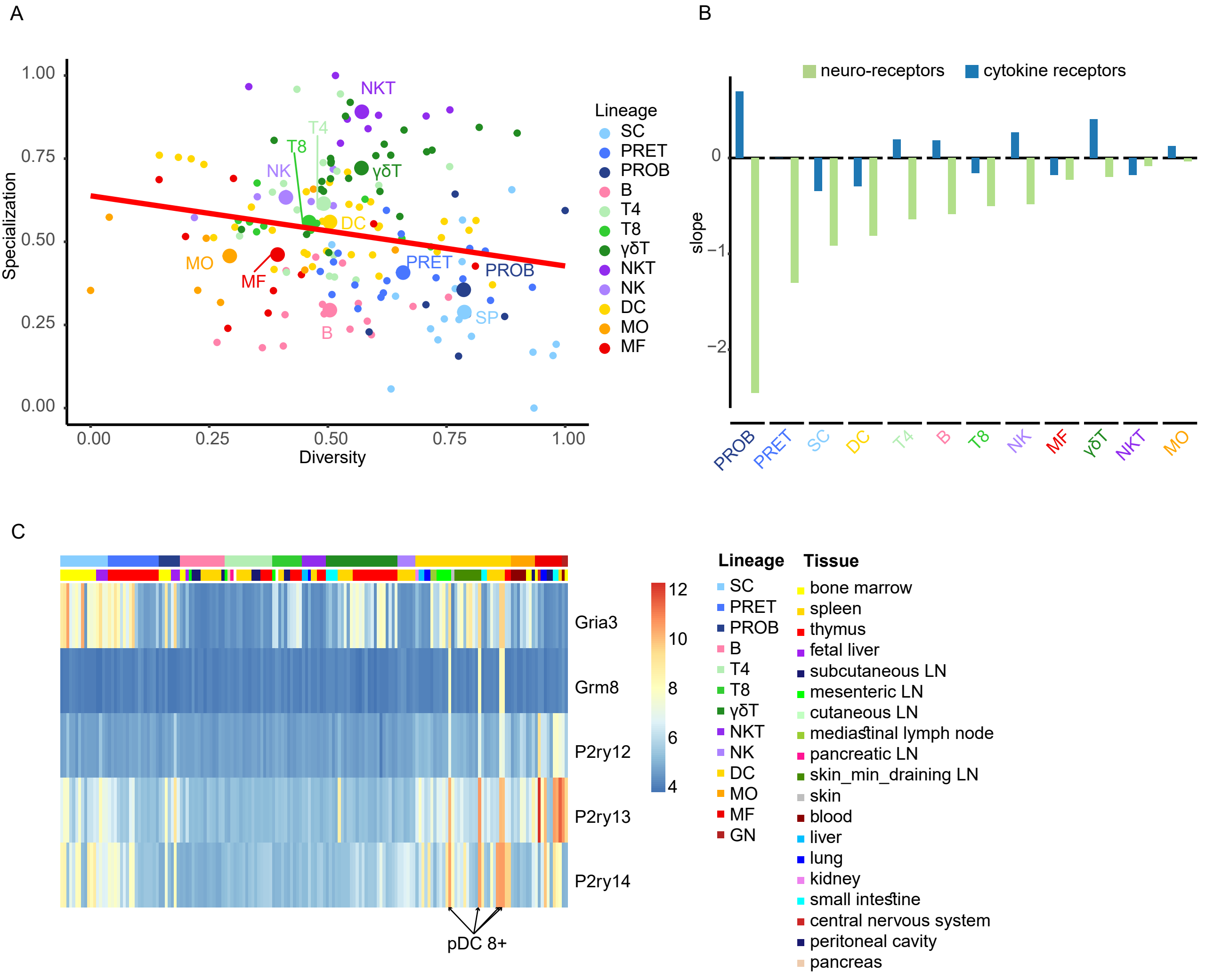


**Figure S4: A negative correlation between cell potential to respond to neural signals and the specificity of signals it can respond to. (A)** Scatter plot of cell diversity versus specialization scores calculated based on cytokine receptors. Scores are scaled ranging 0 (low) to 1 (high). A negative correlation is observed between diversity and specialization of immune cell types (red line, Pearson's correlation coefficient *r= -0.19*). Cells colored by lineage, and lineage centroids are enlarged and labeled. Immune lineages included: stem cells (SC), progenitor B cells (PROB), progenitor T cells (PRET), B cells (B), CD4 T cells (T4), CD8 T cells (T8), gamma-delta T cells (γδT), natural killer T cells (NKT), natural killer (NK), dendritic cells (DC), monocytes (MO) and macrophages (MF). **(B)** Bar plot of the linear regression slope of diversity vs. specialization calculated for each lineage cell types separately. The analysis was done for neuro-receptor and for cytokine receptors separately each represented by different color (green and blue respectively). **(C)** Neuro-receptors with high contribution to the separation of plasmacytoid DC (pDC) from conventional DC (cDC) in PCA represented in Figure 4C. Heatmap of immune cells log2 expression of neuro-receptors with loading score < -0.15 in PC2 (Glutamate receptors- Gria3, Grm8, and purinergic receptors- P2ry12, P2ry13 and P2y14). Immune cells are ordered by lineage and tissue which are annotated by color. pDC are highlighted at the bottom of the heatmap.

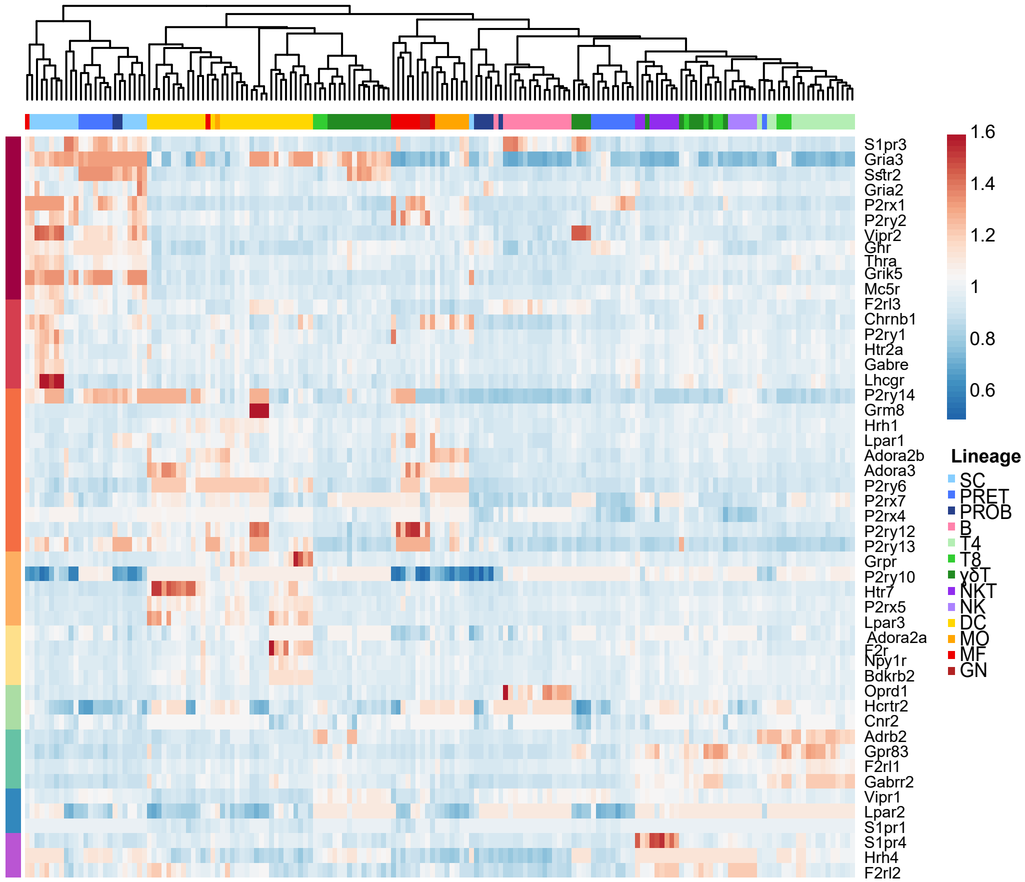


**Figure S5 - Neuro receptors are co-expressed in a lineage specific manner.** Hierarchical clustering of all immune cell types by the 122 neuro-receptors assigned to modules by WGCNA. Genes were ordered by the modules annotated on the left and the expression data was scaled by row. Top color bar indicates the cell type lineage.

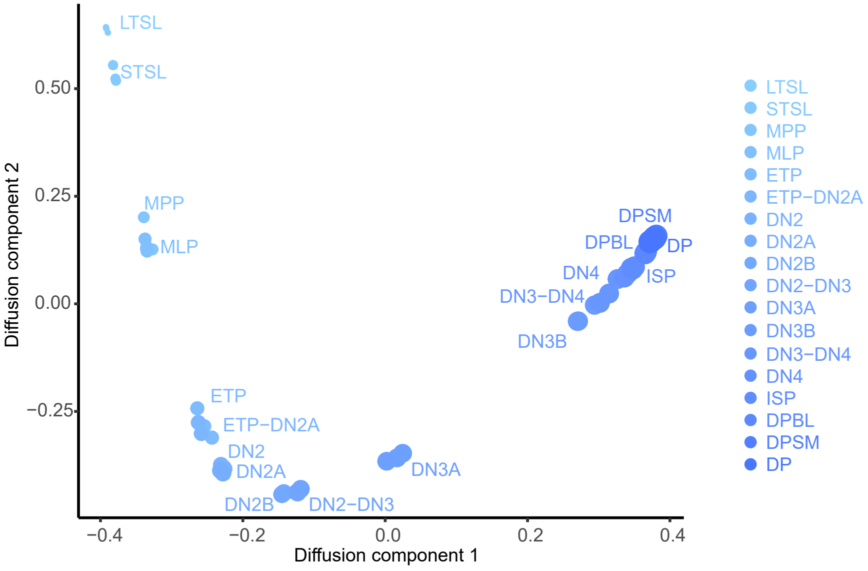


**Figure S6: Neuro-receptor expression profile recapitulates progenitor T cells cellular differentiation.** Diffusion map applied on gene expression of 41 neuro-receptors differentially expressed (one-way ANOVA with *p-value <0.01*) between 15 progenitor T cell types (in replicates) from bone marrow and thymus, creating a trajectory of T cells ordered by the exact stages of differentiation (right) described by gradient color.

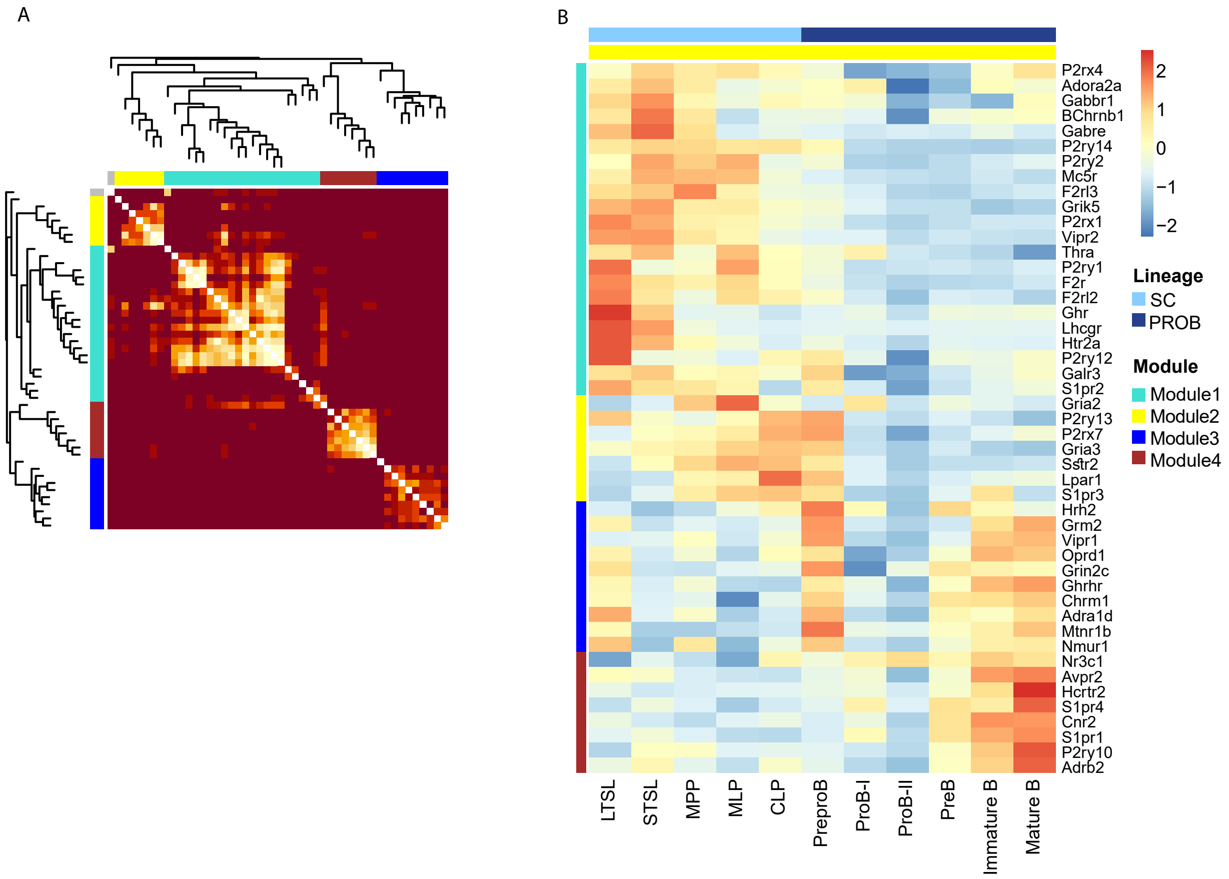


**Figure S7: Neuro-receptors function together to regulate human B cells development.** Neuro-receptors clustered to 4 modules along the B cells developmental trajectory by WGCNA. **(A)** Heatmap plot of the Topological Overlap Matrix (TOM). Darker red color represents low overlap and lighter color represents higher overlap. Blocks of light colors along the diagonal correspond to the identified modules. The gene dendrogram and the assigned module colors are shown along the axes. **(B)** Gene expression heatmap of neuro-receptors assigned to modules and are co-expressed along B cells developmental trajectory. Neuro-receptors are ordered by modules and expression is scaled by row. Cell types are ordered by their differentiation stages and annotated by lineage: stem cells (SC) and progenitor B cells (PROB).

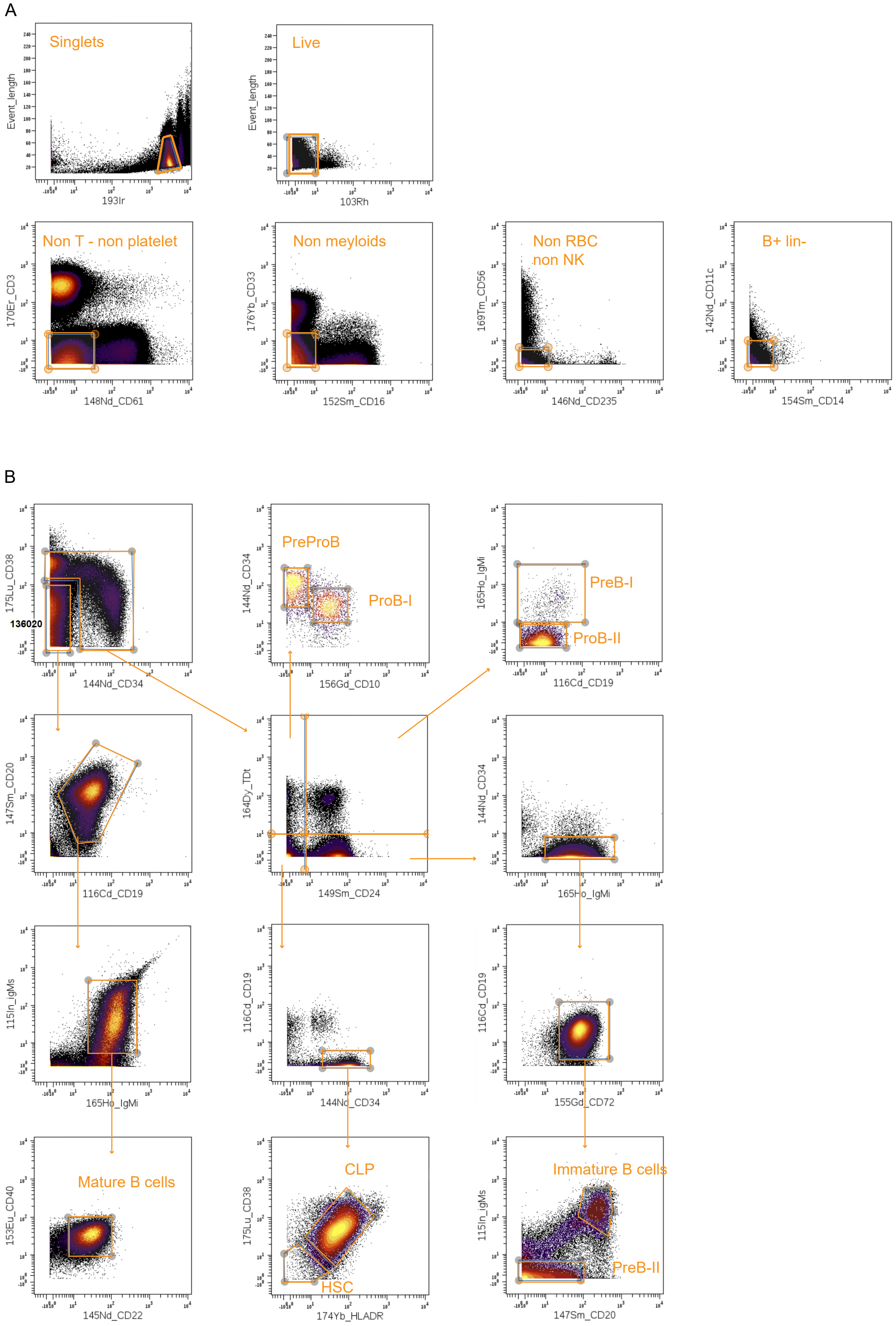


**Figure S8**: **CyTOF manual gating scheme. (A)** Gating strategy for lineage-negative B cells (B+ Lin- cells). Mass cytometry results from a representative healthy donor (3) bone marrow is shown. **(B)** Gating strategy to identify 9 subpopulations of B progenitor cell types among B+ Lin- cells from (A). Hematopoietic stem cells (HSC), common lymphoid progenitor (CLP).
