## Supplementary Table for "Systematic mapping of neural signals in the periphery reveals a rich and specific alphabet for immune cell communication"

**Table S1: Immune cell type.** List of immune cell subsets sorted and characterized by ImmGen with respective name, lineage, and tissue. Scores of cells diversity and specialization were calculated based on Shannon's information theory and scaled ranging 0 - 1.

| Cell subset | Lineage | Tissue | Number of replicates | Specialization scaled | Diversity scaled |
|---|---|---|---|---|---|
| B.FRE.BM | B | bone marrow | 3 | 0.49 | 0.53 |
| B.FRF.BM | B | bone marrow | 3 | 0.54 | 0.30 |
| B.FRE.FL | B | fetal liver | 3 | 0.50 | 0.53 |
| B.FO.MLN | B | mesenteric LN | 3 | 0.56 | 0.30 |
| B.FO.PC | B | peritoneal cavity | 3 | 0.53 | 0.37 |
| B1A.PC | B | peritoneal cavity | 3 | 0.55 | 0.29 |
| B1B.PC | B | peritoneal cavity | 3 | 0.60 | 0.18 |
| B.FO.SP | B | spleen | 3 | 0.63 | 0.06 |
| B.GC.SP | B | spleen | 3 | 0.19 | 0.58 |
| B.MZ.SP | B | spleen | 3 | 0.65 | 0.12 |
| B.T1.SP | B | spleen | 3 | 0.60 | 0.25 |
| B.T2.SP | B | spleen | 3 | 0.63 | 0.16 |
| B.T3.SP | B | spleen | 3 | 0.62 | 0.17 |
| B1A.SP | B | spleen | 6 | 0.59 | 0.23 |
| B.FO.LN | B | subcutaneous LN | 2 | 0.52 | 0.35 |
| DC.103_min_11B_plus_F4/80LO.KD | DC | kidney | 3 | 0.46 | 0.33 |
| DC.103_min_11B_plus_.LV | DC | liver | 3 | 0.49 | 0.50 |
| DC.103_plus_11B_min_.LV | DC | liver | 2 | 0.46 | 0.50 |
| DC.103_min_11B_plus_24_plus_.LU | DC | lung | 2 | 0.44 | 0.43 |
| DC.103_plus_11B_min_.LU | DC | lung | 5 | 0.58 | 0.24 |
| DC.103_min_11B_plus_.LULN | DC | mediastinal lymph node | 3 | 0.63 | 0.44 |
| DC.103_plus_11B_min_.LULN | DC | mediastinal lymph node | 3 | 0.29 | 0.64 |
| DC.4_plus_.MLN | DC | mesenteric LN | 3 | 0.36 | 0.70 |
| DC.8_min_4_min_11B_min_.MLN | DC | mesenteric LN | 3 | 0.53 | 0.61 |
| DC.8_min_4_min_11B_plus_.MLN | DC | mesenteric LN | 3 | 0.49 | 0.63 |
| DC.8_plus_.MLN | DC | mesenteric LN | 3 | 0.53 | 0.44 |
| DC.PDC.8_plus_.MLN | DC | mesenteric LN | 2 | 0.86 | 0.27 |
| DC.LC.SK | DC | skin | 2 | 0.38 | 0.27 |
| DC.4_plus_.SLN | DC | skin_min_draining LN | 3 | 0.24 | 0.79 |

| | | | | | |
|---|---|---|---|---|---|
| DC.8_min_4_min_11B_min_.SLN | DC | skin_min_draining LN | 3 | 0.60 | 0.46 |
| DC.8_min_4_min_11B_plus_.SLN | DC | skin_min_draining LN | 3 | 0.56 | 0.57 |
| DC.8_plus_.SLN | DC | skin_min_draining LN | 3 | 0.47 | 0.50 |
| DC.IIHILANG_min_103_min_11B_plus_.SLN | DC | skin_min_draining LN | 3 | 0.58 | 0.52 |
| DC.IIHILANG_min_103_min_11BLO.SLN | DC | skin_min_draining LN | 3 | 0.48 | 0.65 |
| DC.IIHILANG_plus_103_min_11B_plus_.SLN | DC | skin_min_draining LN | 3 | 0.22 | 0.62 |
| DC.IIHILANG_plus_103_plus_11BLO.SLN | DC | skin_min_draining LN | 3 | 0.33 | 0.52 |
| DC.PDC.8_plus_.SLN | DC | skin_min_draining LN | 3 | 0.90 | 0.24 |
| DC.103_plus_11B_min_.SI | DC | small intestine | 4 | 0.46 | 0.49 |
| DC.103_plus_11B_plus_.SI | DC | small intestine | 4 | 0.29 | 0.64 |
| DC.4_plus_.SP | DC | spleen | 8 | 0.38 | 0.57 |
| DC.8_min_4_min_11B_min_.SP | DC | spleen | 3 | 0.57 | 0.62 |
| DC.8_min_4_min_11B_plus_.SP | DC | spleen | 3 | 0.28 | 0.80 |
| DC.8_plus_.SP | DC | spleen | 8 | 0.64 | 0.31 |
| DC.PDC.8_min_.SP | DC | spleen | 3 | 0.97 | 0.10 |
| DC.PDC.8_plus_.SP | DC | spleen | 3 | 1.00 | 0.09 |
| DC.8_min_.TH | DC | thymus | 3 | 0.82 | 0.00 |
| DC.8_plus_.TH | DC | thymus | 3 | 0.78 | 0.10 |
| TGD.VG5_min_.ACT.IEL | GDT | small intestine | 3 | 0.27 | 0.42 |
| TGD.VG5_min_.IEL | GDT | small intestine | 3 | 0.30 | 0.50 |
| TGD.VG5_plus_.ACT.IEL | GDT | small intestine | 3 | 0.36 | 0.23 |
| TGD.VG5_plus_.IEL | GDT | small intestine | 3 | 0.25 | 0.33 |
| TGD.SP | GDT | spleen | 3 | 0.58 | 0.59 |
| TGD.VG2_min_.ACT.SP | GDT | spleen | 3 | 0.51 | 0.51 |
| TGD.VG2_min_.SP | GDT | spleen | 3 | 0.68 | 0.41 |
| TGD.VG2_plus_.ACT.SP | GDT | spleen | 3 | 0.49 | 0.43 |
| TGD.VG2_plus_.SP | GDT | spleen | 3 | 0.80 | 0.37 |
| TGD.TH | GDT | thymus | 3 | 0.61 | 0.50 |
| TGD.VD1_plus_24AHI.TH | GDT | thymus | 2 | 0.74 | 0.37 |
| TGD.VG1_plus_VD6_min_.TH | GDT | thymus | 3 | 0.70 | 0.37 |
| TGD.VG1_plus_VD6_min_24AHI.TH | GDT | thymus | 3 | 0.78 | 0.42 |
| TGD.VG1_plus_VD6_min_24ALO.TH | GDT | thymus | 2 | 0.46 | 0.68 |

| | | | | | |
|---|---|---|---|---|---|
| TGD.VG1_plus_VD6_plus_.TH | GDT | thymus | 3 | 0.51 | 0.59 |
| TGD.VG1_plus_VD6_plus_24AHI.TH | GDT | thymus | 3 | 0.63 | 0.46 |
| TGD.VG1_plus_VD6_plus_24ALO.TH | GDT | thymus | 3 | 0.45 | 0.35 |
| TGD.VG2_min_24AHI.TH | GDT | thymus | 2 | 0.67 | 0.43 |
| TGD.VG2_plus_24AHI.E17.TH | GDT | thymus | 3 | 0.74 | 0.24 |
| TGD.VG2_plus_24AHI.TH | GDT | thymus | 3 | 0.65 | 0.52 |
| TGD.VG2_plus_24ALO.TH | GDT | thymus | 2 | 0.52 | 0.19 |
| TGD.VG3_plus_24AHI.E17.TH | GDT | thymus | 3 | 0.43 | 0.71 |
| TGD.VG3_plus_24ALO.E17.TH | GDT | thymus | 3 | 0.43 | 0.66 |
| TGD.VG5_plus_24AHI.TH | GDT | thymus | 2 | 0.69 | 0.40 |
| GN.BL | GN | blood | 3 | | |
| GN.BM | GN | bone marrow | 4 | | |
| MF.BM | MF | bone marrow | 3 | 0.52 | 0.48 |
| MF.MICROGLIA.CNS | MF | central nervous system | 3 | 0.45 | 0.07 |
| MF.103_min_11B_plus_.LU | MF | lung | 2 | 0.38 | 0.66 |
| MF.LU | MF | lung | 3 | 0.49 | 0.01 |
| MF.II_min_480HI.PC | MF | peritoneal cavity | 3 | 0.43 | 0.06 |
| MF.II_plus_480LO.PC | MF | peritoneal cavity | 3 | 0.28 | 0.38 |
| MF.103_min_11B_plus_.SI | MF | small intestine | 7 | 0.59 | 0.05 |
| MF.11CLOSER.SI | MF | small intestine | 5 | 0.74 | 0.02 |
| MF.RP.SP | MF | spleen | 3 | 0.30 | 0.08 |
| MO.6C_min_II_min_.BL | MO | blood | 2 | 0.15 | 0.32 |
| MO.6C_min_II_plus_.BL | MO | blood | 2 | 0.22 | 0.74 |
| MO.6C_min_IIINT.BL | MO | blood | 5 | 0.29 | 0.15 |
| MO.6C_plus_II_min_.BL | MO | blood | 3 | 0.40 | 0.30 |
| MO.6C_plus_II_plus_.BL | MO | blood | 2 | 0.29 | 0.54 |
| MO.6C_min_II_min_.BM | MO | bone marrow | 3 | 0.30 | 0.38 |
| MO.6C_plus_II_min_.BM | MO | bone marrow | 3 | 0.33 | 0.58 |
| MO.6C_plus_II_min_.LN | MO | subcutaneous LN | 3 | 0.61 | 0.46 |
| NK.49CI_min_.SP | NK | spleen | 3 | 0.40 | 0.54 |
| NK.49CI_plus_.SP | NK | spleen | 3 | 0.33 | 0.51 |
| NK.49H_min_.SP | NK | spleen | 3 | 0.44 | 0.32 |
| NK.49H_plus_.SP | NK | spleen | 3 | 0.52 | 0.21 |
| NK.B2M_min_.SP | NK | spleen | 3 | 0.40 | 0.38 |
| NK.SP | NK | spleen | 3 | 0.33 | 0.53 |

| | | | | | |
|---|---|---|---|---|---|
| NKT.4_min_.LV | NKT | liver | 4 | 0.31 | 0.43 |
| NKT.4_plus_.LV | NKT | liver | 4 | 0.37 | 0.21 |
| NKT.4_plus_.LU | NKT | lung | 2 | 0.40 | 0.77 |
| NKT.4_min_.SP | NKT | spleen | 3 | 0.40 | 0.37 |
| NKT.4_plus_.SP | NKT | spleen | 3 | 0.29 | 0.47 |
| NKT.44_min_NK1.1_min_.TH | NKT | thymus | 2 | 0.28 | 0.63 |
| NKT.44_plus_NK1.1_min_.TH | NKT | thymus | 3 | 0.45 | 0.37 |
| NKT.44_plus_NK1.1_plus_.TH | NKT | thymus | 3 | 0.25 | 0.48 |
| PRET.DN2.TH | PRET | thymus | 3 | 0.74 | 0.39 |
| PRET.DN2_min_3.TH | PRET | thymus | 2 | 0.59 | 0.49 |
| PRET.DN2A.TH | PRET | thymus | 2 | 0.90 | 0.22 |
| PRET.DN2B.TH | PRET | thymus | 2 | 0.73 | 0.43 |
| PRET.DN3_min_4.TH | PRET | thymus | 3 | 0.24 | 0.85 |
| PRET.DN3A.TH | PRET | thymus | 3 | 0.48 | 0.55 |
| PRET.DN3B.TH | PRET | thymus | 3 | 0.30 | 0.73 |
| PRET.ETP.TH | PRET | thymus | 3 | 0.77 | 0.45 |
| PRET.ETP_min_2A.TH | PRET | thymus | 2 | 0.80 | 0.43 |
| T.DN4.TH | PRET | thymus | 3 | 0.38 | 0.68 |
| T.DP.TH | PRET | thymus | 3 | 0.06 | 1.00 |
| T.DP69_plus_.TH | PRET | thymus | 3 | 0.08 | 0.87 |
| T.DPBL.MHC_min_.TH | PRET | thymus | 3 | 0.19 | 0.77 |
| T.DPBL.TH | PRET | thymus | 3 | 0.19 | 0.70 |
| T.DPSM.MHC_min_.TH | PRET | thymus | 2 | 0.23 | 0.75 |
| T.DPSM.TH | PRET | thymus | 3 | 0.14 | 0.81 |
| T.ISP.TH | PRET | thymus | 3 | 0.23 | 0.76 |
| PREB.FRC.BM | PROB | bone marrow | 3 | 0.00 | 0.79 |
| PREB.FRD.BM | PROB | bone marrow | 3 | 0.36 | 0.68 |
| PROB.FRA.BM | PROB | bone marrow | 4 | 0.53 | 0.66 |
| PROB.FRBC.BM | PROB | bone marrow | 3 | 0.26 | 0.76 |
| PREB.FRD.FL | PROB | fetal liver | 3 | 0.42 | 0.62 |
| PROB.FRA.FL | PROB | fetal liver | 3 | 0.87 | 0.49 |
| PROB.FRBC.FL | PROB | fetal liver | 3 | 0.20 | 0.78 |
| MLP.BM | SC | bone marrow | 4 | 0.75 | 0.38 |
| PROB.CLP.BM | SC | bone marrow | 3 | 0.70 | 0.46 |
| SC.CDP.BM | SC | bone marrow | 3 | 0.60 | 0.45 |
| SC.CMP.BM | SC | bone marrow | 5 | 0.49 | 0.59 |
| SC.GMP.BM | SC | bone marrow | 3 | 0.30 | 0.67 |
| SC.LT34F.BM | SC | bone marrow | 3 | 0.85 | 0.14 |
| SC.LTSL.BM | SC | bone marrow | 3 | 0.56 | 0.46 |
| SC.MDP.BM | SC | bone marrow | 3 | 0.44 | 0.56 |
| SC.MEP.BM | SC | bone marrow | 2 | 0.13 | 0.54 |
| SC.MPP34F.BM | SC | bone marrow | 2 | 0.66 | 0.49 |
| SC.ST34F.BM | SC | bone marrow | 2 | 0.76 | 0.27 |
| SC.STSL.BM | SC | bone marrow | 3 | 0.78 | 0.26 |

| | | | | | |
|---|---|---|---|---|---|
| MLP.FL | SC | fetal liver | 3 | 0.71 | 0.46 |
| PROB.CLP.FL | SC | fetal liver | 3 | 0.83 | 0.46 |
| SC.LTSL.FL | SC | fetal liver | 3 | 0.52 | 0.67 |
| SC.STSL.FL | SC | fetal liver | 2 | 0.69 | 0.58 |
| BEC.SLN | STROMAL | cutaneous LN | 5 | | |
| FRC.SLN | STROMAL | cutaneous LN | 8 | | |
| LEC.SLN | STROMAL | cutaneous LN | 4 | | |
| ST.31_min_38_min_44_min_.SLN | STROMAL | cutaneous LN | 3 | | |
| BEC.MLN | STROMAL | mesenteric LN | 8 | | |
| FRC.MLN | STROMAL | mesenteric LN | 8 | | |
| LEC.MLN | STROMAL | mesenteric LN | 6 | | |
| FI.SK | STROMAL | skin | 4 | | |
| EP.MECHI.TH | STROMAL | thymus | 3 | | |
| FI.MTS15_plus_.TH | STROMAL | thymus | 3 | | |
| T.4NVE.MLN | T4 | mesenteric LN | 3 | 0.42242 | 0.444597 |
| T.4.PA.BDC | T4 | pancreas | 2 | 0.452691 | 0.530901 |
| T.4.PLN.BDC | T4 | pancreatic LN | 3 | 0.472688 | 0.462909 |
| T.4NVE.PP | T4 | Peyers patches | 2 | 0.4634 | 0.456729 |
| T.4FP3_min_.SP | T4 | spleen | 3 | 0.455193 | 0.455277 |
| T.4FP3_plus_25_plus_.SP | T4 | spleen | 2 | 0.487653 | 0.344724 |
| T.4MEM.SP | T4 | spleen | 3 | 0.528433 | 0.231046 |
| T.4MEM44H62L.SP | T4 | spleen | 3 | 0.508694 | 0.307594 |
| T.4NVE.SP | T4 | spleen | 4 | 0.618419 | 0.149302 |
| T.4.LN.BDC | T4 | subcutaneous LN | 3 | 0.47601 | 0.421816 |
| T.4MEM.LN | T4 | subcutaneous LN | 3 | 0.401147 | 0.51772 |
| T.4NVE.LN | T4 | subcutaneous LN | 3 | 0.505147 | 0.321094 |
| T.4_plus_8INT.TH | T4 | thymus | 3 | 0.149015 | 0.867134 |
| T.4SP24_min_.TH | T4 | thymus | 3 | 0.521321 | 0.332076 |
| T.4SP24INT.TH | T4 | thymus | 3 | 0.248668 | 0.620182 |
| T.4SP69_plus_.TH | T4 | thymus | 3 | 0.338686 | 0.573603 |
| T.8NVE.MLN | T8 | mesenteric LN | 3 | 0.6351 | 0.361907 |
| T.8NVE.PP | T8 | Peyers patches | 3 | 0.459753 | 0.651884 |
| T.8MEM.SP | T8 | spleen | 3 | 0.512168 | 0.290136 |
| T.8NVE.SP | T8 | spleen | 3 | 0.438447 | 0.608508 |
| T.8MEM.LN | T8 | subcutaneous LN | 3 | 0.32619 | 0.577145 |
| T.8NVE.LN | T8 | subcutaneous LN | 3 | 0.634883 | 0.268065 |
| T.4INT8_plus_.TH | T8 | thymus | 3 | 0.386263 | 0.697543 |
| T.8SP24_min_.TH | T8 | thymus | 3 | 0.660501 | 0.316932 |
| T.8SP24INT.TH | T8 | thymus | 3 | 0.613968 | 0.52432 |
| T.8SP69_plus_.TH | T8 | thymus | 3 | 0.506817 | 0.640663 |

**Table S2: Neuronal signals receptors and ligands.** Listing all neuro-receptors and neural ligands from KEGG pathway labeled 'Neuroactive ligand-receptor interaction - Mus musculus', reviewing their neuronal signal name and type (neurotransmitter (NT), neuropeptide (NP), neurohormone (NH)), the expression threshold determined for each gene separately, the expression status (Expressed / Not expressed) and the number of lineages the gene is expressed in.

| Symbol | Type | Signal name | Class | Expression threshold | Expressed in Dataset | lineages expressed |
|---|---|---|---|---|---|---|
| Chrm1 | Receptor | Acetylcholine | NT | 6.9 | TRUE | 14 |
| Chrm2 | Receptor | Acetylcholine | NT | 6.9 | FALSE | 0 |
| Chrm3 | Receptor | Acetylcholine | NT | 6 | TRUE | 6 |
| Chrm4 | Receptor | Acetylcholine | NT | 6.1 | TRUE | 10 |
| Chrm5 | Receptor | Acetylcholine | NT | 6.9 | FALSE | 0 |
| Chrna1 | Receptor | Acetylcholine | NT | 6.9 | FALSE | 0 |
| Chrna10 | Receptor | Acetylcholine | NT | 6.9 | FALSE | 0 |
| Chrna2 | Receptor | Acetylcholine | NT | 6.9 | FALSE | 0 |
| Chrna3 | Receptor | Acetylcholine | NT | 6.9 | FALSE | 0 |
| Chrna4 | Receptor | Acetylcholine | NT | 6.9 | TRUE | 1 |
| Chrna5 | Receptor | Acetylcholine | NT | 6.9 | TRUE | 1 |
| Chrna6 | Receptor | Acetylcholine | NT | 6.9 | FALSE | 0 |
| Chrna7 | Receptor | Acetylcholine | NT | 6.9 | FALSE | 0 |
| Chrna9 | Receptor | Acetylcholine | NT | 6.5 | TRUE | 1 |
| Chrnb1 | Receptor | Acetylcholine | NT | 6.8 | TRUE | 8 |
| Chrnb2 | Receptor | Acetylcholine | NT | 6.9 | FALSE | 0 |
| Chrnb3 | Receptor | Acetylcholine | NT | 5.5 | TRUE | 1 |
| Chrnb4 | Receptor | Acetylcholine | NT | 6.9 | TRUE | 3 |
| Chrnd | Receptor | Acetylcholine | NT | 6.9 | FALSE | 0 |
| Chrne | Receptor | Acetylcholine | NT | 6.9 | TRUE | 1 |
| Chrng | Receptor | Acetylcholine | NT | 6.1 | TRUE | 14 |
| Chat | Ligand | Acetylcholine | NT | 6.9 | FALSE | 0 |
| Adora1 | Receptor | Adenosine | NT | 6.9 | FALSE | 0 |
| Adora2a | Receptor | Adenosine | NT | 6.9 | TRUE | 13 |
| Adora2b | Receptor | Adenosine | NT | 6.2 | TRUE | 7 |
| Adora3 | Receptor | Adenosine | NT | 6.6 | TRUE | 5 |
| Cnr1 | Receptor | Anandamide | NT | 6.9 | FALSE | 0 |
| Cnr2 | Receptor | Anandamide | NT | 6.2 | TRUE | 14 |
| Trpv1 | Receptor | Anandamide | NT | 6.9 | FALSE | 0 |
| Napepld | Ligand | Anandamide | NT | 6.1 | TRUE | 4 |
| Agtr1a | Receptor | Angiotensin | NH | 6.9 | TRUE | 1 |
| Agtr1b | Receptor | Angiotensin | NH | 6.9 | TRUE | 7 |
| Agtr2 | Receptor | Angiotensin | NH | 5.8 | TRUE | 1 |
| Mas1 | Receptor | Angiotensin | NH | 6.9 | FALSE | 0 |
| Agt | Ligand | Angiotensin | NH | 6.9 | TRUE | 8 |
| Aplnr | Receptor | Apelin | NP | 6.3 | TRUE | 4 |
| Apln | Ligand | Apelin | NP | 6.9 | TRUE | 2 |

| | | | | | | |
|---|---|---|---|---|---|---|
| Avpr1a | Receptor | arginine vasopressin | NH | 6.9 | TRUE | 1 |
| Avpr1b | Receptor | arginine vasopressin | NH | 6.9 | FALSE | 0 |
| Avpr2 | Receptor | arginine vasopressin | NH | 6.4 | TRUE | 13 |
| Avp | Ligand | arginine vasopressin | NH | 6.9 | TRUE | 7 |
| P2rx1 | Receptor | ATP | NT | 6 | TRUE | 8 |
| P2rx2 | Receptor | ATP | NT | 6.3 | TRUE | 2 |
| P2rx3 | Receptor | ATP | NT | 6.9 | TRUE | 1 |
| P2rx4 | Receptor | ATP | NT | 6.9 | TRUE | 12 |
| P2rx5 | Receptor | ATP | NT | 6.8 | TRUE | 2 |
| P2rx6 | Receptor | ATP | NT | 6.9 | TRUE | 1 |
| P2rx7 | Receptor | ATP | NT | 6.9 | TRUE | 12 |
| P2ry1 | Receptor | ATP | NT | 6.4 | TRUE | 11 |
| P2ry10 | Receptor | ATP | NT | 6.9 | TRUE | 9 |
| P2ry12 | Receptor | ATP | NT | 5.1 | TRUE | 11 |
| P2ry13 | Receptor | ATP | NT | 5.9 | TRUE | 10 |
| P2ry14 | Receptor | ATP | NT | 6.3 | TRUE | 9 |
| P2ry2 | Receptor | ATP | NT | 6.4 | TRUE | 4 |
| P2ry2 | Receptor | ATP | NT | 6.4 | TRUE | 4 |
| P2ry4 | Receptor | ATP | NT | 6.9 | FALSE | 0 |
| P2ry6 | Receptor | ATP | NT | 6.9 | TRUE | 5 |
| Kng1 | Ligand | Bradykinin | NT | 5.8 | TRUE | 1 |
| Bdkrb1 | Receptor | Bradykinin | NP | 6.1 | TRUE | 13 |
| Bdkrb2 | Receptor | Bradykinin | NP | 6.9 | TRUE | 10 |
| Calcr | Receptor | Calcitonin | NP | 6.9 | FALSE | 0 |
| Calca | Ligand | Calcitonin | NP | 6.9 | FALSE | 0 |
| Cckar | Receptor | Cholecystokinin | NP | 6.9 | FALSE | 0 |
| Cckbr | Receptor | Cholecystokinin | NP | 6.9 | FALSE | 0 |
| Cck | Ligand | Cholecystokinin | NP | 6.9 | FALSE | 0 |
| Crhr1 | Receptor | Corticotropin Releasing neuro-hormone | NH | 6.9 | TRUE | 1 |
| Crhr2 | Receptor | Corticotropin Releasing neuro-hormone | NH | 6.9 | FALSE | 0 |
| Crh | Ligand | Corticotropin Releasing neuro-hormone | NH | 6.9 | FALSE | 0 |
| Nr3c1 | Receptor | Cortisol | NH | 6.9 | TRUE | 14 |
| Cyp11b1 | Ligand | Cortisol | NH | 6.9 | FALSE | 0 |
| Drd1a | Receptor | Dopamine | NT | 6.9 | TRUE | 1 |
| Drd2 | Receptor | Dopamine | NT | 6.9 | FALSE | 0 |
| Drd3 | Receptor | Dopamine | NT | 6.9 | FALSE | 0 |
| Drd4 | Receptor | Dopamine | NT | 6.9 | TRUE | 14 |
| Drd5 | Receptor | Dopamine | NT | 6.9 | FALSE | 0 |
| Ddc | Ligand | Dopamine | NT | 6.9 | TRUE | 1 |
| Oprd1 | Receptor | Dynorphins | NP | 6.9 | TRUE | 13 |

| | | | | | | |
|---|---|---|---|---|---|---|
| Oprk1 | Receptor | Dynorphins | NP | 6.9 | FALSE | 0 |
| Pdyn | Ligand | Dynorphins | NP | 6.9 | FALSE | 0 |
| Oprm1 | Receptor | Endorphin | NP | 6.9 | FALSE | 0 |
| Pomc | Ligand | Endorphin | NP | 6.9 | TRUE | 1 |
| Ednra | Receptor | Endothelin | NP | 6.7 | TRUE | 3 |
| Ednrb | Receptor | Endothelin | NP | 6.3 | TRUE | 2 |
| Edn1 | Ligand | Endothelin | NP | 6.4 | TRUE | 3 |
| Oprd1 | Receptor | Enkephalin | NP | 6.9 | TRUE | 13 |
| Penk | Ligand | Enkephalin | NP | 5.6 | TRUE | 7 |
| Adra1a | Receptor | Epinephrine | NT | 6.9 | TRUE | 1 |
| Adra1b | Receptor | Epinephrine | NT | 6.9 | FALSE | 0 |
| Adra1d | Receptor | Epinephrine | NT | 6.9 | TRUE | 14 |
| Adra2a | Receptor | Epinephrine | NT | 6.9 | TRUE | 1 |
| Adra2b | Receptor | Epinephrine | NT | 6.9 | TRUE | 1 |
| Adra2c | Receptor | Epinephrine | NT | 6.9 | FALSE | 0 |
| Adrb1 | Receptor | Epinephrine | NT | 6.9 | TRUE | 12 |
| Adrb2 | Receptor | Epinephrine | NT | 6.9 | TRUE | 14 |
| Adrb3 | Receptor | Epinephrine | NT | 6.9 | FALSE | 0 |
| Pnmt | Ligand | Epinephrine | NT | 6.9 | FALSE | 0 |
| Fshr | Receptor | FSH | NH | 6.9 | FALSE | 0 |
| Fshb | Ligand | FSH | NH | 6.9 | FALSE | 0 |
| Gabbr1 | Receptor | GABA | NT | 6.9 | TRUE | 13 |
| Gabbr2 | Receptor | GABA | NT | 6.9 | FALSE | 0 |
| Gabra1 | Receptor | GABA | NT | 6.9 | FALSE | 0 |
| Gabra2 | Receptor | GABA | NT | 6.9 | FALSE | 0 |
| Gabra3 | Receptor | GABA | NT | 6 | TRUE | 1 |
| Gabra4 | Receptor | GABA | NT | 6.9 | FALSE | 0 |
| Gabra5 | Receptor | GABA | NT | 5.6 | TRUE | 1 |
| Gabra6 | Receptor | GABA | NT | 6.9 | FALSE | 0 |
| Gabrb1 | Receptor | GABA | NT | 6.9 | FALSE | 0 |
| Gabrb2 | Receptor | GABA | NT | 4.8 | TRUE | 1 |
| Gabrb3 | Receptor | GABA | NT | 6.9 | FALSE | 0 |
| Gabrd | Receptor | GABA | NT | 6.9 | FALSE | 0 |
| Gabre | Receptor | GABA | NT | 5.2 | TRUE | 2 |
| Gabrg1 | Receptor | GABA | NT | 6.9 | FALSE | 0 |
| Gabrg2 | Receptor | GABA | NT | 6.9 | FALSE | 0 |
| Gabrg3 | Receptor | GABA | NT | 6.3 | TRUE | 1 |
| Gabrp | Receptor | GABA | NT | 4.8 | TRUE | 13 |
| Gabrq | Receptor | GABA | NT | 5.1 | TRUE | 2 |
| Gabrr1 | Receptor | GABA | NT | 6.9 | FALSE | 0 |
| Gabrr2 | Receptor | GABA | NT | 6.3 | TRUE | 10 |
| Gabrr3 | Receptor | GABA | NT | 6.9 | FALSE | 0 |
| Gad1 | Ligand | GABA | NT | 6.9 | TRUE | 2 |
| Galr1 | Receptor | Galanin | NP | 6.9 | FALSE | 0 |
| Galr2 | Receptor | Galanin | NP | 6.9 | FALSE | 0 |
| Galr3 | Receptor | Galanin | NP | 6.9 | TRUE | 14 |

| | | | | | | |
|---|---|---|---|---|---|---|
| Gal | Ligand | Galanin | NP | 6.9 | TRUE | 1 |
| Grpr | Receptor | Gastrin Releasing neuro-peptide | NP | 5.6 | TRUE | 2 |
| Grp | Ligand | Gastrin Releasing neuro-peptide | NP | 6.9 | TRUE | 1 |
| Ghsr | Receptor | Ghrelin | NH | 6.8 | TRUE | 13 |
| Ghrl | Ligand | Ghrelin | NH | 6.9 | TRUE | 14 |
| Glp1r | Receptor | Glucagon-like neuro-peptide-1 | NP | 6.6 | TRUE | 2 |
| Gcg | Ligand | Glucagon-like neuro-peptide-1 | NP | 6.9 | TRUE | 1 |
| Glp2r | Receptor | Glucagon-like neuro-peptide-2 | NP | 6.9 | FALSE | 0 |
| Gcg | Ligand | Glucagon-like neuro-peptide-2 | NP | 6.9 | TRUE | 1 |
| Gria1 | Receptor | Glutamate | NT | 6.9 | FALSE | 0 |
| Gria2 | Receptor | Glutamate | NT | 4.8 | TRUE | 2 |
| Gria3 | Receptor | Glutamate | NT | 5.9 | TRUE | 8 |
| Gria4 | Receptor | Glutamate | NT | 5 | TRUE | 1 |
| Grid1 | Receptor | Glutamate | NT | 6.9 | FALSE | 0 |
| Grid2 | Receptor | Glutamate | NT | 6.9 | FALSE | 0 |
| Grik1 | Receptor | Glutamate | NT | 6.9 | FALSE | 0 |
| Grik2 | Receptor | Glutamate | NT | 6.9 | FALSE | 0 |
| Grik3 | Receptor | Glutamate | NT | 6.9 | FALSE | 0 |
| Grik4 | Receptor | Glutamate | NT | 6.9 | FALSE | 0 |
| Grik5 | Receptor | Glutamate | NT | 6.6 | TRUE | 9 |
| Grin1 | Receptor | Glutamate | NT | 6.9 | FALSE | 0 |
| Grin2a | Receptor | Glutamate | NT | 6.9 | FALSE | 0 |
| Grin2b | Receptor | Glutamate | NT | 6.9 | FALSE | 0 |
| Grin2c | Receptor | Glutamate | NT | 6 | TRUE | 13 |
| Grin2d | Receptor | Glutamate | NT | 6.9 | TRUE | 1 |
| Grin3a | Receptor | Glutamate | NT | 6.9 | FALSE | 0 |
| Grin3b | Receptor | Glutamate | NT | 6.9 | TRUE | 4 |
| Grm1 | Receptor | Glutamate | NT | 6.9 | FALSE | 0 |
| Grm2 | Receptor | Glutamate | NT | 6.9 | TRUE | 14 |
| Grm3 | Receptor | Glutamate | NT | 6.9 | FALSE | 0 |
| Grm4 | Receptor | Glutamate | NT | 6.1 | TRUE | 12 |
| Grm5 | Receptor | Glutamate | NT | 6.9 | FALSE | 0 |
| Grm6 | Receptor | Glutamate | NT | 6.9 | TRUE | 1 |
| Grm7 | Receptor | Glutamate | NT | 6.9 | FALSE | 0 |
| Grm8 | Receptor | Glutamate | NT | 6.9 | TRUE | 1 |
| Gls | Ligand | Glutamate | NT | 6.9 | TRUE | 14 |
| Glra1 | Receptor | Glycine | NT | 6.9 | FALSE | 0 |
| Glra2 | Receptor | Glycine | NT | 6.9 | FALSE | 0 |
| Glra3 | Receptor | Glycine | NT | 6.9 | FALSE | 0 |
| Glra4 | Receptor | Glycine | NT | 6.9 | FALSE | 0 |

| | | | | | | |
|---|---|---|---|---|---|---|
| Grin2a | Receptor | Glycine | NT | 6.9 | FALSE | 0 |
| Gnrh1 | Ligand | Gonadotropin-releasing neuro-hormone | NH | 6.9 | TRUE | 5 |
| Gnrhr | Receptor | Gonadotropin-releasing neuro-hormone | NP | 6.9 | FALSE | 0 |
| Ghr | Receptor | Growth neuro-hormone | NH | 5.7 | TRUE | 2 |
| Gh | Ligand | Growth neuro-hormone | NH | 6.9 | TRUE | 2 |
| Ghrhr | Receptor | Growth neuro-hormone Releasing neuro-hormone | NH | 6.9 | TRUE | 14 |
| Ghrh | Ligand | Growth neuro-hormone Releasing neuro-hormone | NH | 5.8 | TRUE | 12 |
| Hrh1 | Receptor | Histamine | NT | 6.9 | TRUE | 11 |
| Hrh2 | Receptor | Histamine | NT | 6.9 | TRUE | 13 |
| Hrh3 | Receptor | Histamine | NT | 6.9 | FALSE | 0 |
| Hrh4 | Receptor | Histamine | NT | 5.2 | TRUE | 2 |
| Hdc | Ligand | Histamine | NT | 6.8 | TRUE | 7 |
| Kiss1r | Receptor | Kiss1 neuro-peptide | NP | 6.5 | TRUE | 2 |
| Kiss1 | Ligand | Kiss1 neuro-peptide | NP | 6.9 | TRUE | 11 |
| Lepr | Receptor | Leptin | NT | 5.1 | TRUE | 3 |
| Lep | Ligand | Leptin | NT | 6.9 | FALSE | 0 |
| Lhcgr | Receptor | LH | NH | 5.6 | TRUE | 2 |
| Lhb | Ligand | LH | NH | 6.9 | TRUE | 9 |
| Lpar1 | Receptor | Lysophosphatidic acid | NT | 6.8 | TRUE | 5 |
| Lpar2 | Receptor | Lysophosphatidic acid | NT | 6.2 | TRUE | 12 |
| Lpar3 | Receptor | Lysophosphatidic acid | NT | 6.3 | TRUE | 1 |
| Lpar4 | Receptor | Lysophosphatidic acid | NT | 6 | TRUE | 1 |
| Mchr1 | Receptor | Melanin concentrating neuro-hormone | NH | 6.9 | FALSE | 0 |
| Pmch | Ligand | Melanin concentrating neuro-hormone | NH | 5.3 | TRUE | 9 |
| Mc1r | Receptor | Melanocortin | NP | 6.3 | TRUE | 10 |
| Mc2r | Receptor | Melanocortin | NP | 5.4 | TRUE | 1 |
| Mc3r | Receptor | Melanocortin | NP | 6.9 | FALSE | 0 |
| Mc4r | Receptor | Melanocortin | NP | 6.9 | FALSE | 0 |
| Mc5r | Receptor | Melanocortin | NP | 6.5 | TRUE | 3 |
| Pomc | Ligand | Melanocortin | NP | 6.9 | TRUE | 1 |
| Gpr50 | Receptor | Melatonin | NH | 6.9 | TRUE | 1 |
| Mtnr1a | Receptor | Melatonin | NH | 6.9 | FALSE | 0 |
| Mtnr1b | Receptor | Melatonin | NH | 5.5 | TRUE | 13 |
| Aanat | Ligand | Melatonin | NH | 5.4 | TRUE | 14 |
| Brs3 | Receptor | Neuromedin B | NP | 6.9 | FALSE | 0 |
| Nmbr | Receptor | Neuromedin B | NP | 6.9 | FALSE | 0 |
| Nmb | Ligand | Neuromedin B | NP | 6.7 | TRUE | 7 |
| Nmur1 | Receptor | Neuromedin U | NP | 6.3 | TRUE | 13 |

| | | | | | | |
|---|---|---|---|---|---|---|
| Nmur2 | Receptor | Neuromedin U | NP | 6.9 | FALSE | 0 |
| Nmu | Ligand | Neuromedin U | NP | 6.9 | FALSE | 0 |
| Npbwr1 | Receptor | Neuroneuro-peptide B | NP | 6.9 | TRUE | 3 |
| Npb | Ligand | Neuroneuro-peptide B | NP | 6.9 | TRUE | 4 |
| Mas1 | Receptor | Neuroneuro-peptide FF | NP | 6.9 | FALSE | 0 |
| Npffr1 | Receptor | Neuroneuro-peptide FF | NP | 6.9 | FALSE | 0 |
| Npffr2 | Receptor | Neuroneuro-peptide FF | NP | 6.9 | FALSE | 0 |
| Npff | Ligand | Neuroneuro-peptide FF | NP | 6.9 | TRUE | 14 |
| Npbwr1 | Receptor | Neuroneuro-peptide W | NP | 6.9 | TRUE | 3 |
| Npw | Ligand | Neuroneuro-peptide W | NP | 6.9 | TRUE | 14 |
| Gpr83 | Receptor | Neuroneuro-peptide Y | NP | 6.8 | TRUE | 4 |
| Npy1r | Receptor | Neuroneuro-peptide Y | NP | 5.6 | TRUE | 2 |
| Npy2r | Receptor | Neuroneuro-peptide Y | NP | 6.9 | FALSE | 0 |
| Npy5r | Receptor | Neuroneuro-peptide Y | NP | 6.9 | FALSE | 0 |
| Npy6r | Receptor | Neuroneuro-peptide Y | NP | 6.9 | FALSE | 0 |
| Npy | Ligand | Neuroneuro-peptide Y | NP | 6.9 | TRUE | 5 |
| Ntsr1 | Receptor | Neurotensin | NP | 6.9 | FALSE | 0 |
| Ntsr2 | Receptor | Neurotensin | NP | 6.9 | TRUE | 11 |
| Nts | Ligand | Neurotensin | NP | 5.5 | TRUE | 1 |
| Oprl1 | Receptor | Nociceptin | NP | 6.9 | FALSE | 0 |
| Pnoc | Ligand | Nociceptin | NP | 6.9 | TRUE | 10 |
| Adra1a | Receptor | Norepinephrine | NT | 6.9 | TRUE | 1 |
| Adra1b | Receptor | Norepinephrine | NT | 6.9 | FALSE | 0 |
| Adra1d | Receptor | Norepinephrine | NT | 6.9 | TRUE | 14 |
| Adra2a | Receptor | Norepinephrine | NT | 6.9 | TRUE | 1 |
| Adra2b | Receptor | Norepinephrine | NT | 6.9 | TRUE | 1 |
| Adra2c | Receptor | Norepinephrine | NT | 6.9 | FALSE | 0 |
| Adrb1 | Receptor | Norepinephrine | NT | 6.9 | TRUE | 12 |
| Adrb2 | Receptor | Norepinephrine | NT | 6.9 | TRUE | 14 |
| Adrb3 | Receptor | Norepinephrine | NT | 6.9 | FALSE | 0 |
| Dbh | Ligand | Norepinephrine | NT | 6.9 | FALSE | 0 |
| Hcrtr1 | Receptor | Orexin | NP | 6.9 | FALSE | 0 |
| Hcrtr2 | Receptor | Orexin | NP | 5.5 | TRUE | 1 |
| Hcrt | Ligand | Orexin | NP | 6.9 | FALSE | 0 |
| Avpr1a | Receptor | Oxytocin | NH | 6.9 | TRUE | 1 |
| Oxtr | Receptor | Oxytocin | NH | 6.9 | FALSE | 0 |
| Pam | Ligand | Oxytocin | NH | 6.9 | TRUE | 12 |
| Adcyap1r1 | Receptor | PACAP | NP | 6.9 | TRUE | 1 |
| Adcyap1 | Ligand | PACAP | NP | 6.9 | FALSE | 0 |
| Pth1r | Receptor | Parathyroid neuro-hormone | NH | 6.5 | TRUE | 1 |
| Pth2r | Receptor | Parathyroid neuro-hormone | NH | 6.9 | FALSE | 0 |
| Pth | Ligand | Parathyroid neuro-hormone | NH | 6.9 | FALSE | 0 |
| Prlr | Receptor | Prolactin | NP | 4.7 | TRUE | 4 |

| | | | | | | |
|---|---|---|---|---|---|---|
| Prl | Ligand | Prolactin | NP | 6.9 | FALSE | 0 |
| Prlhr | Receptor | Prolactin releasing neuro-peptide | NP | 6.9 | FALSE | 0 |
| Prlh | Ligand | Prolactin releasing neuro-peptide | NP | 6.9 | TRUE | 14 |
| F2r | Receptor | Proteinase activated like | NP | 6.9 | TRUE | 11 |
| F2rl1 | Receptor | Proteinase activated like | NP | 6.3 | TRUE | 7 |
| F2rl2 | Receptor | Proteinase activated like | NP | 6.9 | TRUE | 8 |
| F2rl3 | Receptor | Proteinase activated like | NP | 6.6 | TRUE | 11 |
| Pard3 | Receptor | Proteinase activated like | NP | 6.9 | TRUE | 2 |
| F2 | Ligand | Proteinase activated like | NP | 6 | TRUE | 2 |
| S1pr1 | Receptor | S1P | NT | 6.5 | TRUE | 14 |
| S1pr2 | Receptor | S1P | NT | 6.9 | TRUE | 14 |
| S1pr3 | Receptor | S1P | NT | 6.3 | TRUE | 8 |
| S1pr4 | Receptor | S1P | NT | 6.9 | TRUE | 14 |
| S1pr5 | Receptor | S1P | NT | 6.7 | TRUE | 4 |
| Sphk1 | Ligand | S1P | NT | 6.9 | TRUE | 5 |
| Sctr | Receptor | Secretin | NH | 6.9 | FALSE | 0 |
| Sct | Ligand | Secretin | NH | 6.9 | TRUE | 1 |
| Htr1a | Receptor | Serotonin | NT | 6.9 | FALSE | 0 |
| Htr1b | Receptor | Serotonin | NT | 6.9 | TRUE | 1 |
| Htr1d | Receptor | Serotonin | NT | 6.9 | FALSE | 0 |
| Htr1f | Receptor | Serotonin | NT | 6.9 | FALSE | 0 |
| Htr2a | Receptor | Serotonin | NT | 6 | TRUE | 4 |
| Htr2b | Receptor | Serotonin | NT | 6.9 | FALSE | 0 |
| Htr2c | Receptor | Serotonin | NT | 6.9 | TRUE | 1 |
| Htr3a | Receptor | Serotonin | NT | 6.9 | FALSE | 0 |
| Htr3b | Receptor | Serotonin | NT | 6.9 | FALSE | 0 |
| Htr4 | Receptor | Serotonin | NT | 6.9 | FALSE | 0 |
| Htr5a | Receptor | Serotonin | NT | 6.9 | FALSE | 0 |
| Htr5b | Receptor | Serotonin | NT | 6.9 | FALSE | 0 |
| Htr6 | Receptor | Serotonin | NT | 6.9 | FALSE | 0 |
| Htr7 | Receptor | Serotonin | NT | 6.4 | TRUE | 2 |
| Ddc | Ligand | Serotonin | NT | 6.9 | TRUE | 1 |
| Sstr1 | Receptor | Somatostatin | NH | 6.9 | FALSE | 0 |
| Sstr2 | Receptor | Somatostatin | NH | 6.6 | TRUE | 4 |
| Sstr3 | Receptor | Somatostatin | NH | 6.9 | FALSE | 0 |
| Sstr4 | Receptor | Somatostatin | NH | 6.8 | TRUE | 5 |
| Sstr5 | Receptor | Somatostatin | NH | 6.9 | FALSE | 0 |
| Sst | Ligand | Somatostatin | NH | 6.9 | TRUE | 1 |

| | | | | | | |
|---|---|---|---|---|---|---|
| Tacr1 | Receptor | Substance P | NP | 6.9 | TRUE | 1 |
| Tac1 | Ligand | Substance P | NP | 5 | TRUE | 1 |
| Tacr2 | Receptor | Tachykinin | NP | 6.9 | FALSE | 0 |
| Tacr3 | Receptor | Tachykinin | NP | 6.9 | FALSE | 0 |
| Tac2 | Ligand | Tachykinin | NP | 6.9 | TRUE | 1 |
| Trhr | Receptor | Thyrotropin Releasing neuro-hormone | NH | 4.7 | FALSE | 0 |
| Trhr2 | Receptor | Thyrotropin Releasing neuro-hormone | NH | 6.9 | FALSE | 0 |
| Trh | Ligand | Thyrotropin Releasing neuro-hormone | NH | 6.9 | FALSE | 0 |
| Thra | Receptor | Thyroxine | NH | 6.7 | TRUE | 13 |
| Tshb | Ligand | Thyroxine | NH | 6.9 | FALSE | 0 |
| Thrb | Receptor | Thyroxine | NT | 6.9 | FALSE | 0 |
| Tshr | Receptor | TSH | NH | 5 | TRUE | 2 |
| Tshb | Ligand | TSH | NH | 6.9 | FALSE | 0 |
| Taar1 | Receptor | Tyramine | NT | 6.9 | FALSE | 0 |
| Taar2 | Receptor | Tyramine | NT | 6.9 | FALSE | 0 |
| Taar3 | Receptor | Tyramine | NT | 4.9 | TRUE | 11 |
| Taar4 | Receptor | Tyramine | NT | 6.9 | FALSE | 0 |
| Taar5 | Receptor | Tyramine | NT | 6.9 | FALSE | 0 |
| Taar6 | Receptor | Tyramine | NT | 6.9 | FALSE | 0 |
| Taar7a | Receptor | Tyramine | NT | 6.9 | FALSE | 0 |
| Taar7b | Receptor | Tyramine | NT | 6.9 | FALSE | 0 |
| Taar7d | Receptor | Tyramine | NT | 6.9 | FALSE | 0 |
| Taar7e | Receptor | Tyramine | NT | 6.9 | FALSE | 0 |
| Taar7f | Receptor | Tyramine | NT | 6.9 | FALSE | 0 |
| Taar8a | Receptor | Tyramine | NT | 6.9 | FALSE | 0 |
| Taar8b | Receptor | Tyramine | NT | 6.9 | FALSE | 0 |
| Taar8c | Receptor | Tyramine | NT | 6.9 | FALSE | 0 |
| Taar9 | Receptor | Tyramine | NT | 6.9 | FALSE | 0 |
| Ddc | Ligand | Tyramine | NT | 6.9 | TRUE | 1 |
| Uts2r | Receptor | Urotensin II | NP | 6.9 | FALSE | 0 |
| Uts2 | Ligand | Urotensin II | NP | 6.9 | FALSE | 0 |
| Vipr1 | Receptor | Vasoactive Intestinal neuro-peptide | NP | 6.1 | TRUE | 14 |
| Vipr2 | Receptor | Vasoactive Intestinal neuro-peptide | NP | 6 | TRUE | 6 |
| Vip | Ligand | Vasoactive Intestinal neuro-peptide | NP | 6.9 | FALSE | 0 |

**Table S3 – Neural ligands defined by synthesis enzyme or precursor gene.** Neural ligands which cannot be detected via gene expression, such as chemicals, are listed and estimated by the gene expression levels of their precursors or enzyme in synthesis.

| **Neural signal** | **Symbol** | **Gene name** |
|---|---|---|
| Cortisol | Cyp11b1 | 11-beta-hydroxylase |
| Melatonin | Aanat | serotonin N-acetyltransferase |
| Oxytocin | Pam | peptidylglycine alpha-amidating monooxygenase |
| Dynorphins | Pdyn | prodynorphin (Opioid) |
| Endorphin | Pomc | proopiomelanocortin |
| Enkephalin | Penk | proenkephalin |
| Glucagon-like neuro-peptide-1 | Gcg | Preproglucagon |
| Glucagon-like neuro-peptide-2 | Gcg | Preproglucagon |
| Melanocortin | Pomc | proopiomelanocortin |
| Orexin | Hcrt | Hypocretin Neuroneuro-peptide Precursor |
| Substance P | Tac1 | Tryptophan hydroxylase |
| Tachykinin | Tac2 | Tryptophan hydroxylase |
| Acetylcholine | Chat | Choline acetyltransferase |
| Anandamide | Napepld | N-acetylphosphatidylethanolamine-hydrolysing phospholipase D |
| Bradykinin | Kng1 | kininogen |
| Dopamine | Ddc | Dopa Decarboxylase |
| Epinephrine | Pnmt | phenylethanolamine N-methyltransferase (PNMT) |
| GABA | Gad1 | glutamate decarboxylase |
| Glutamate | Gls | Glutaminase |
| Histamine | Hdc | L-histidine decarboxylase |
| Norepinephrine | Dbh | dopamine beta-hydroxylase |
| S1P | Sphk1 | sphingosine kinase |
| Serotonin | Ddc | Dopa Decarboxylase |

**Table S4 – Cytokine receptors list.** Listing cytokine receptors symbol, description, threshold expression determined for each gene separately and the expression status (Expressed / Not expressed).

| Symbol | Description | Expression threshold | Expressed |
|---|---|---|---|
| Acvr1 | activin A receptor, type 1 | 5.9 | TRUE |
| Acvr1b | activin A receptor, type 1B | 6.9 | TRUE |
| Acvr1c | activin A receptor, type IC | 6.9 | FALSE |
| Acvr2a | activin receptor IIA | 6.9 | TRUE |
| Acvr2b | activin receptor IIB | 6.6 | TRUE |
| Acvrl1 | activin A receptor, type II-like 1 | 6.9 | TRUE |
| Amhr2 | anti-Mullerian hormone type 2 receptor | 5.3 | TRUE |
| Bmpr1a | bone morphogenetic protein receptor, type 1A | 5.3 | TRUE |
| Bmpr2 | bone morphogenetic protein receptor, type II (serine/threonine kinase) | 6.9 | TRUE |
| Ccr1 | chemokine (C-C motif) receptor 1 | 5.8 | TRUE |
| Ccr10 | chemokine (C-C motif) receptor 10 | 6.9 | TRUE |
| Ccr11 | chemokine (C-C motif) receptor 11 | 6.9 | FALSE |
| Ccr2 | chemokine (C-C motif) receptor 2 | 6.4 | TRUE |
| Ccr3 | chemokine (C-C motif) receptor 3 | 5.2 | TRUE |
| Ccr4 | chemokine (C-C motif) receptor 4 | 6.1 | TRUE |
| Ccr5 | chemokine (C-C motif) receptor 5 | 6.6 | TRUE |
| Ccr6 | chemokine (C-C motif) receptor 6 | 6.5 | TRUE |
| Ccr7 | chemokine (C-C motif) receptor 7 | 6.9 | TRUE |
| Ccr8 | chemokine (C-C motif) receptor 8 | 5.4 | TRUE |
| Ccr9 | chemokine (C-C motif) receptor 9 | 6.6 | TRUE |
| Cd27 | CD27 antigen | 6.9 | TRUE |
| Cd4 | CD4 antigen | 6.9 | TRUE |
| Cd40 | CD40 antigen | 6.9 | TRUE |
| Cntfr | ciliary neurotrophic factor receptor | 6.9 | TRUE |
| Crlf2 | cytokine receptor-like factor 2 | 6.9 | TRUE |
| Csf1r | colony stimulating factor 1 receptor | 6.9 | TRUE |
| Csf2ra | colony stimulating factor 2 receptor, alpha, low-affinity (granulocyte-macrophage) | 6.9 | TRUE |
| Csf2rb | colony stimulating factor 2 receptor, beta, low-affinity (granulocyte-macrophage) | 6.8 | TRUE |
| Csf2rb2 | colony stimulating factor 2 receptor, beta 2, low-affinity (granulocyte-macrophage) | 6.9 | TRUE |
| Csf3r | colony stimulating factor 3 receptor (granulocyte) | 6.2 | TRUE |
| Cx3cr1 | chemokine (C-X3-C motif) receptor 1 | 6.9 | TRUE |
| Cxcr1 | chemokine (C-X-C motif) receptor 1 | 6.4 | TRUE |
| Cxcr2 | chemokine (C-X-C motif) receptor 2 | 5 | TRUE |
| Cxcr3 | chemokine (C-X-C motif) receptor 3 | 6.9 | TRUE |
| Cxcr4 | chemokine (C-X-C motif) receptor 4 | 6.9 | TRUE |
| Cxcr5 | chemokine (C-X-C motif) receptor 5 | 5.5 | TRUE |
| Cxcr6 | chemokine (C-X-C motif) receptor 6 | 6.2 | TRUE |
| Cxcr7 | chemokine (C-X-C motif) receptor 7 | 5.8 | TRUE |

| | | | |
|---|---|---|---|
| Eda2r | ectodysplasin A2 receptor | 5.6 | FALSE |
| Edar | ectodysplasin-A receptor | 6.5 | TRUE |
| Epor | erythropoietin receptor | 6.9 | TRUE |
| Fas | Fas (TNF receptor superfamily member 6) | 6.3 | TRUE |
| Ghr | growth hormone receptor | 5.7 | TRUE |
| Gm13305 | predicted gene 13305 | 6.9 | FALSE |
| Ifnar1 | interferon (alpha and beta) receptor 1 | 6.9 | TRUE |
| Ifnar2 | interferon (alpha and beta) receptor 2 | 6.9 | TRUE |
| Ifngr1 | interferon gamma receptor 1 | 6.9 | TRUE |
| Ifngr2 | interferon gamma receptor 2 | 6.9 | TRUE |
| Ifnlr1 | interferon lambda receptor 1 | 6.9 | FALSE |
| Il10ra | interleukin 10 receptor, alpha | 6.9 | TRUE |
| Il10rb | interleukin 10 receptor, beta | 6.9 | TRUE |
| Il11ra1 | interleukin 11 receptor, alpha chain 1 | 6.9 | TRUE |
| Il11ra2 | interleukin 11 receptor, alpha chain 2 | 6.9 | TRUE |
| Il12rb1 | interleukin 12 receptor, beta 1 | 6.9 | TRUE |
| Il13ra1 | interleukin 13 receptor, alpha 1 | 6.3 | TRUE |
| Il13ra2 | interleukin 13 receptor, alpha 2 | 4.6 | FALSE |
| Il15ra | interleukin 15 receptor, alpha chain | 6.9 | TRUE |
| Il17ra | interleukin 17 receptor A | 6.9 | TRUE |
| Il17rb | interleukin 17 receptor B | 6.5 | TRUE |
| Il17rc | interleukin 17 receptor C | 6.9 | TRUE |
| Il17re | interleukin 17 receptor E | 6.8 | TRUE |
| Il18r1 | interleukin 18 receptor 1 | 5.8 | TRUE |
| Il18rap | interleukin 18 receptor accessory protein | 6.8 | TRUE |
| Il1r1 | interleukin 1 receptor, type I | 5.7 | TRUE |
| Il1r2 | interleukin 1 receptor, type II | 6.5 | TRUE |
| Il1rap | interleukin 1 receptor accessory protein | 6.9 | TRUE |
| Il1rl1 | interleukin 1 receptor-like 1 | 5.6 | TRUE |
| Il1rl2 | interleukin 1 receptor-like 2 | 5.2 | TRUE |
| Il20ra | interleukin 20 receptor, alpha | 5.5 | TRUE |
| Il20rb | interleukin 20 receptor beta | 6.9 | TRUE |
| Il21r | interleukin 21 receptor | 6.9 | TRUE |
| Il22ra1 | interleukin 22 receptor, alpha 1 | 6.9 | FALSE |
| Il23r | interleukin 23 receptor | 5.8 | TRUE |
| Il27ra | interleukin 27 receptor, alpha | 6.9 | TRUE |
| Il2ra | interleukin 2 receptor, alpha chain | 6.1 | TRUE |
| Il2rb | interleukin 2 receptor, beta chain | 6.9 | TRUE |
| Il2rg | interleukin 2 receptor, gamma chain | 6.9 | TRUE |
| Il3ra | interleukin 3 receptor, alpha chain | 6.9 | TRUE |
| Il31ra | interleukin 31 receptor A | 5.2 | TRUE |
| Il4ra | interleukin 4 receptor, alpha | 6.9 | TRUE |
| Il5ra | interleukin 5 receptor, alpha | 5.2 | TRUE |
| Il6ra | interleukin 6 receptor, alpha | 6.9 | TRUE |
| Il6st | interleukin 6 signal transducer | 6.9 | TRUE |
| Il7r | interleukin 7 receptor | 6 | TRUE |

| | | | |
|---|---|---|---|
| Il9r | interleukin 9 receptor | 6.9 | TRUE |
| Lepr | leptin receptor | 6.6 | TRUE |
| Lifr | LIF receptor alpha | 6.4 | TRUE |
| Ltbr | lymphotoxin B receptor | 6.9 | TRUE |
| Mpl | myeloproliferative leukemia virus oncogene | 6.9 | TRUE |
| Ngfr | nerve growth factor receptor (TNFR superfamily, member 16) | 6.9 | TRUE |
| Osmr | oncostatin M receptor | 6.5 | FALSE |
| Prlr | prolactin receptor | 4.8 | TRUE |
| Relt | RELT tumor necrosis factor receptor | 6.9 | TRUE |
| Tgfbr1 | transforming growth factor, beta receptor I | 6.9 | TRUE |
| Tgfbr2 | transforming growth factor, beta receptor II | 6.9 | TRUE |
| Tnfrsf10b | tumor necrosis factor receptor superfamily, member 10b | 6.1 | TRUE |
| Tnfrsf11a | tumor necrosis factor receptor superfamily, member 11a, NFKB activator | 6.8 | TRUE |
| Tnfrsf11b | tumor necrosis factor receptor superfamily, member 11b (osteoprotegerin) | 6 | TRUE |
| Tnfrsf12a | tumor necrosis factor receptor superfamily, member 12a | 6.9 | TRUE |
| Tnfrsf13b | tumor necrosis factor receptor superfamily, member 13b | 6.9 | TRUE |
| Tnfrsf13c | tumor necrosis factor receptor superfamily, member 13c | 6.9 | TRUE |
| Tnfrsf14 | tumor necrosis factor receptor superfamily, member 14 (herpesvirus entry mediator) | 6.9 | TRUE |
| Tnfrsf17 | tumor necrosis factor receptor superfamily, member 17 | 6.3 | TRUE |
| Tnfrsf18 | tumor necrosis factor receptor superfamily, member 18 | 6.7 | TRUE |
| Tnfrsf19 | tumor necrosis factor receptor superfamily, member 19 | 6.8 | TRUE |
| Tnfrsf1a | tumor necrosis factor receptor superfamily, member 1a | 6.9 | TRUE |
| Tnfrsf1b | tumor necrosis factor receptor superfamily, member 1b | 6.9 | TRUE |
| Tnfrsf21 | tumor necrosis factor receptor superfamily, member 21 | 6.9 | TRUE |
| Tnfrsf25 | tumor necrosis factor receptor superfamily, member 25 | 6.5 | TRUE |
| Tnfrsf4 | tumor necrosis factor receptor superfamily, member 4 | 6.9 | TRUE |
| Tnfrsf8 | tumor necrosis factor receptor superfamily, member 8 | 6.9 | TRUE |
| Tnfrsf9 | tumor necrosis factor receptor superfamily, member 9 | 6.8 | TRUE |
| Xcr1 | chemokine (C motif) receptor 1 | 6.9 | TRUE |

**Table S5 – Neuro-receptors involved in neuro-immune communication network in the intestine.** List of neuro-receptors expressed on immune cells included in intestinal neuro-immune communication network. MM – muscularis residing macrophages, LpM - lamina propria residing macrophages.

| Neuro-receptor | Neural signal | Immune cell | Log2(expression) |
|---|---|---|---|
| Chrm1 | Acetylcholine | ɣδT Vγ5- | |
| Chrm4 | Acetylcholine | ɣδT Vγ5- | |
| Chrna1 | Acetylcholine | ɣδT Vγ5- | |
| Chrnb1 | Acetylcholine | ɣδT Vγ5- | |
| Chrng | Acetylcholine | ɣδT Vγ5- | |
| Chrm1 | Acetylcholine | ɣδT Vγ5+ | |
| Chrna1 | Acetylcholine | ɣδT Vγ5+ | |
| Chrnb1 | Acetylcholine | ɣδT Vγ5+ | |
| Chrng | Acetylcholine | ɣδT Vγ5+ | |
| Ddc | Dopamine | ɣδT Vγ5- | |
| Drd4 | Dopamine | ɣδT Vγ5+ | |
| Ddc | Dopamine | ɣδT Vγ5+ | |
| Gabbr1 | GABA | DC CD103+CD11b- | |
| Gabbr1 | GABA | DC CD103+CD11b+ | |
| Gabbr1 | GABA | ɣδT Vγ5- | |
| Gabrp | GABA | ɣδT Vγ5- | |
| Gad1 | GABA | ɣδT Vγ5- | |
| Gabrp | GABA | ɣδT Vγ5+ | |
| Gad1 | GABA | ɣδT Vγ5+ | |
| Galr3 | Galanin | ɣδT Vγ5- | |
| Galr3 | Galanin | ɣδT Vγ5+ | |
| Ghrl | Ghrelin | DC CD103+CD11b+ | |
| Ghsr | Ghrelin | ɣδT Vγ5- | |
| Ghrl | Ghrelin | ɣδT Vγ5- | |
| Ghsr | Ghrelin | ɣδT Vγ5+ | |
| Ghrl | Ghrelin | MF CD103-CD11b+ (LpM) | |
| Ghrl | Ghrelin | MF CD11c- (MM) | |
| Glp1r | Glucagon-like peptide-1 | ɣδT Vγ5- | |
| Gcg | Glucagon-like peptide-1 | ɣδT Vγ5- | |
| Glp1r | Glucagon-like peptide-1 | ɣδT Vγ5+ | |
| Gcg | Glucagon-like peptide-1 | ɣδT Vγ5+ | |
| Gls | Glutamate | DC CD103+CD11b- | |
| Gls | Glutamate | DC CD103+CD11b+ | |
| Grm2 | Glutamate | ɣδT Vγ5- | |
| Grm4 | Glutamate | ɣδT Vγ5- | |
| Gls | Glutamate | ɣδT Vγ5- | |
| Grm2 | Glutamate | ɣδT Vγ5+ | |
| Grm4 | Glutamate | ɣδT Vγ5+ | |
| Gls | Glutamate | ɣδT Vγ5+ | |
| Gls | Glutamate | MF CD103-CD11b+ (LpM) | |
| Gls | Glutamate | MF CD11c- (MM) | |

| | | |
|---|---|---|
| Adra2b | Norepinephrine | DC CD103+CD11b- |
| Adrb2 | Norepinephrine | DC CD103+CD11b- |
| Adra2b | Norepinephrine | DC CD103+CD11b+ |
| Adrb2 | Norepinephrine | DC CD103+CD11b+ |
| Adra1d | Norepinephrine | ɣδT Vγ5- |
| Adrb1 | Norepinephrine | ɣδT Vγ5- |
| Adra1d | Norepinephrine | ɣδT Vγ5+ |
| Adra2b | Norepinephrine | ɣδT Vγ5+ |
| Adrb1 | Norepinephrine | ɣδT Vγ5+ |
| Adra2b | Norepinephrine | MF CD103-CD11b+ (LpM) |
| Adrb1 | Norepinephrine | MF CD103-CD11b+ (LpM) |
| Adrb2 | Norepinephrine | MF CD103-CD11b+ (LpM) |
| Adra2b | Norepinephrine | MF CD11c- (MM) |
| Adrb2 | Norepinephrine | MF CD11c- (MM) |
| Vipr1 | VIP | ɣδT Vγ5- |
| Vipr2 | VIP | ɣδT Vγ5- |
| Vipr1 | VIP | ɣδT Vγ5+ |
| Vipr2 | VIP | ɣδT Vγ5+ |
| Vipr1 | VIP | MF CD103-CD11b+ (LpM) |

**Table S6 - Enrichment analysis of neuro-receptors assigned to modules correlated to lineage.** List of modules of neuro-receptors constructed using WGCNA. The cells lineage trait correlation to module eigengene was calculated and significant correlations (Bonferroni corrected p-value < 0.05) are listed in Lineages column. For each module, enriched Go terms are listed providing information of Bonferroni correction p-values and the neuro-receptors enriched.

| module | Lineages | genes | Go Term | p.adjust | Genes enriched |
|---|---|---|---|---|---|
| 1 | SC, PreT | P2rx1/ P2ry2/ Gria2/ Gria3/ Grik5/ Mc5r/ F2rl3/ S1pr3/ Sstr2/ Thra/ Vipr2 | | | |
| 2 | SC | Chrnb1/ P2ry1/ Gabre/ Ghr/ Lhcgr/ Htr2a | small molecule metabolic process | 0.298 | Ghr/ Htr2a/ Lhcgr/ P2ry1 |
| | | | positive regulation of carbohydrate metabolic process | 0.298 | Htr2a/ Lhcgr/ P2ry1 |
| | | | organophosphate biosynthetic process | 0.298 | Htr2a/ Lhcgr/ P2ry1 |
| 3 | DC, MF, MO | Adora2b/ Adora3/ P2rx4/ P2rx7/ P2ry12/ P2ry13/ P2ry14/ P2ry6/ Grm8/ Hrh1/ Lpar1 | positive regulation of immune system process | 0.047 | Adora2b/ Adora3/ P2rx4/ P2rx7/ P2ry12 |
| | | | positive regulation of cell migration | 0.033 | Adora2b/ Adora3/ Lpar1/ P2rx4/ P2ry12/ P2ry6 |

| | | | | | |
|---|---|---|---|---|---|
| | | | positive regulation of cellular component movement | 0.033 | Adora2b/ Adora3/ Lpar1/ P2rx4/ P2ry12/ P2ry6 |
| 4 | DC | P2rx5/ P2ry10/ Grpr/ Lpar3/ Htr7 | | | |
| 5 | DC | Adora2a/ Bdkrb2/ Oprd1/ Npy1r | regulation of apoptotic signaling pathway | 0.325 | Adora2a/ Bdkrb2 |
| | | | negative regulation of macromolecule metabolic process | 0.387 | Adora2a/ Bdkrb2/ Oprd1 |
| | | | locomotory behavior | 0.387 | Adora2a/ Npy1r/ Oprd1 |
| 6 | B | Cnr2/ Adrb2/ Hcrtr2 | regulation of response to stress | 0.432 | Adrb2/ Cnr2 |
| | | | negative regulation of inflammatory response | 0.345 | Adrb2/ Cnr2 |
| | | | negative regulation of cell activation | 0.345 | Adrb2/ Cnr2 |
| 7 | Lympho-cytes | Gabrr2/ Gpr83/ F2rl1/ Vipr1 | | | |
| 8 | T4, T8 | Lpar2/ S1pr1/ S1pr4 | | | |
| 9 | NKT, NK | Hrh4/F2r/ F2rl2 | | | |

**Table S7 - Neuro-receptors and cytokine receptors lineage association.** List of significant correlations (Bonferroni corrected p-value < 0.05) between neuro-receptor and cytokine receptors expression profile and lineage cell types binary vector. For each correlated pair of lineage-receptor, information of Pearson's correlation and Benferroni correction p-values is provided.

| Gene symbol | Lineage | Correlation | Adjusted pValue | Receptor type |
|---|---|---|---|---|
| Il5ra | B | 0.90 | 4.47E-60 | cytokine receptor |
| Tnfrsf13c | B | 0.89 | 9.38E-57 | cytokine receptor |
| Cxcr5 | B | 0.85 | 2.71E-45 | cytokine receptor |
| Tnfrsf13b | B | 0.84 | 3.45E-43 | cytokine receptor |
| Ccr6 | B | 0.61 | 1.52E-15 | cytokine receptor |
| Il9r | B | 0.58 | 5.96E-13 | cytokine receptor |
| Cd40 | B | 0.40 | 0.00019 | cytokine receptor |
| Hcrtr2 | B | 0.76 | 5.71E-30 | neuro-receptor |
| Cnr2 | B | 0.57 | 2.21E-12 | neuro-receptor |
| Avpr2 | B | 0.41 | 7.15E-05 | neuro-receptor |
| S1pr1 | B | 0.39 | 0.000349 | neuro-receptor |
| Chrnb1 | B | 0.35 | 0.006144 | neuro-receptor |
| S1pr3 | B | 0.34 | 0.021612 | neuro-receptor |
| Csf2rb2 | DC | 0.78 | 4.00E-32 | cytokine receptor |
| Tnfrsf11a | DC | 0.72 | 1.33E-24 | cytokine receptor |

| | | | | |
|---|---|---|---|---|
| Csf2rb | DC | 0.69 | 1.10E-21 | cytokine receptor |
| Il13ra1 | DC | 0.66 | 5.03E-19 | cytokine receptor |
| Acvr1 | DC | 0.62 | 2.89E-16 | cytokine receptor |
| Cd40 | DC | 0.62 | 5.29E-16 | cytokine receptor |
| Xcr1 | DC | 0.59 | 9.71E-14 | cytokine receptor |
| Acvr2a | DC | 0.57 | 2.36E-12 | cytokine receptor |
| Csf2ra | DC | 0.56 | 3.95E-12 | cytokine receptor |
| Il15ra | DC | 0.56 | 1.10E-11 | cytokine receptor |
| Il10ra | DC | 0.49 | 2.48E-08 | cytokine receptor |
| Bmpr2 | DC | 0.44 | 3.67E-06 | cytokine receptor |
| Ifngr2 | DC | 0.43 | 1.11E-05 | cytokine receptor |
| Tnfrsf4 | DC | 0.43 | 1.92E-05 | cytokine receptor |
| Lifr | DC | 0.42 | 3.73E-05 | cytokine receptor |
| Acvrl1 | DC | 0.42 | 4.66E-05 | cytokine receptor |
| Ccr7 | DC | 0.42 | 4.66E-05 | cytokine receptor |
| Il3ra | DC | 0.41 | 7.99E-05 | cytokine receptor |
| Ccr5 | DC | 0.37 | 0.001685 | cytokine receptor |
| Acvr2b | DC | 0.37 | 0.002408 | cytokine receptor |
| Il1r2 | DC | 0.33 | 0.030315 | cytokine receptor |
| Ltbr | DC | 0.33 | 0.039372 | cytokine receptor |
| Lpar3 | DC | 0.72 | 3.18E-25 | neuro-receptor |
| P2rx5 | DC | 0.69 | 5.69E-22 | neuro-receptor |
| Htr7 | DC | 0.69 | 1.47E-21 | neuro-receptor |
| Hrh1 | DC | 0.58 | 6.15E-13 | neuro-receptor |
| Oprd1 | DC | 0.55 | 2.61E-11 | neuro-receptor |
| P2ry6 | DC | 0.50 | 1.35E-08 | neuro-receptor |
| Grpr | DC | 0.48 | 8.55E-08 | neuro-receptor |
| Adora3 | DC | 0.44 | 4.08E-06 | neuro-receptor |
| P2ry14 | DC | 0.43 | 1.94E-05 | neuro-receptor |
| P2ry10 | DC | 0.40 | 0.000241 | neuro-receptor |
| Npy1r | DC | 0.39 | 0.000455 | neuro-receptor |
| Bdkrb2 | DC | 0.39 | 0.000591 | neuro-receptor |
| Adra2b | DC | 0.38 | 0.001238 | neuro-receptor |
| P2rx4 | DC | 0.34 | 0.01781 | neuro-receptor |
| Grm8 | DC | 0.33 | 0.031299 | neuro-receptor |
| Il12rb1 | GDT | 0.60 | 1.30E-14 | cytokine receptor |
| Il2rb | GDT | 0.53 | 4.19E-10 | cytokine receptor |
| Cxcr6 | GDT | 0.52 | 7.17E-10 | cytokine receptor |
| Ccr8 | GDT | 0.51 | 3.25E-09 | cytokine receptor |
| Ccr10 | GDT | 0.50 | 9.11E-09 | cytokine receptor |
| Ccr9 | GDT | 0.49 | 2.81E-08 | cytokine receptor |
| Ccr4 | GDT | 0.49 | 4.28E-08 | cytokine receptor |
| Il20ra | GDT | 0.47 | 3.25E-07 | cytokine receptor |
| Tnfrsf9 | GDT | 0.44 | 5.95E-06 | cytokine receptor |
| Il27ra | GDT | 0.41 | 9.20E-05 | cytokine receptor |
| Cd27 | GDT | 0.39 | 0.000584 | cytokine receptor |

| | | | | |
|---|---|---|---|---|
| Lpar2 | GDT | 0.47 | 4.67E-07 | neuro-receptor |
| P2rx7 | GDT | 0.36 | 0.003122 | neuro-receptor |
| F2r | GDT | 0.36 | 0.003925 | neuro-receptor |
| Glp1r | GDT | 0.36 | 0.005738 | neuro-receptor |
| Cxcr1 | GN | 0.73 | 4.65E-26 | cytokine receptor |
| Cxcr2 | GN | 0.67 | 7.16E-20 | cytokine receptor |
| Il1rap | GN | 0.58 | 1.90E-13 | cytokine receptor |
| Bmpr1a | GN | 0.46 | 5.91E-07 | cytokine receptor |
| Csf3r | GN | 0.43 | 1.08E-05 | cytokine receptor |
| Ccr1 | GN | 0.40 | 0.000143 | cytokine receptor |
| Il1r2 | GN | 0.35 | 0.007218 | cytokine receptor |
| Tshr | GN | 0.61 | 3.74E-15 | neuro-receptor |
| Kiss1r | GN | 0.59 | 3.58E-14 | neuro-receptor |
| Ednra | GN | 0.58 | 3.26E-13 | neuro-receptor |
| Chrm3 | GN | 0.51 | 3.06E-09 | neuro-receptor |
| Hrh2 | GN | 0.50 | 1.47E-08 | neuro-receptor |
| P2ry2 | GN | 0.34 | 0.015693 | neuro-receptor |
| Csf1r | MF | 0.50 | 7.90E-09 | cytokine receptor |
| Ccr1 | MF | 0.45 | 2.34E-06 | cytokine receptor |
| Tnfrsf21 | MF | 0.43 | 1.49E-05 | cytokine receptor |
| Epor | MF | 0.43 | 2.15E-05 | cytokine receptor |
| Tgfbr1 | MF | 0.40 | 0.000219 | cytokine receptor |
| Ccr3 | MF | 0.38 | 0.00084 | cytokine receptor |
| Ltbr | MF | 0.38 | 0.000868 | cytokine receptor |
| Csf2ra | MF | 0.38 | 0.001253 | cytokine receptor |
| Acvr1 | MF | 0.37 | 0.001434 | cytokine receptor |
| Il11ra2 | MF | 0.36 | 0.003454 | cytokine receptor |
| Il11ra1 | MF | 0.35 | 0.007224 | cytokine receptor |
| Csf3r | MF | 0.34 | 0.015542 | cytokine receptor |
| Tnfrsf11a | MF | 0.34 | 0.017279 | cytokine receptor |
| Ccr5 | MF | 0.32 | 0.046283 | cytokine receptor |
| Il13ra1 | MF | 0.32 | 0.048112 | cytokine receptor |
| Tnfrsf12a | MF | 0.32 | 0.048269 | cytokine receptor |
| Cx3cr1 | MF | 0.32 | 0.049972 | cytokine receptor |
| P2ry12 | MF | 0.57 | 7.60E-13 | neuro-receptor |
| Lpar1 | MF | 0.50 | 9.16E-09 | neuro-receptor |
| P2rx4 | MF | 0.49 | 2.08E-08 | neuro-receptor |
| Ednrb | MF | 0.47 | 4.82E-07 | neuro-receptor |
| P2ry13 | MF | 0.46 | 7.29E-07 | neuro-receptor |
| P2ry2 | MF | 0.38 | 0.001132 | neuro-receptor |
| Adora3 | MF | 0.37 | 0.002799 | neuro-receptor |
| Adrb1 | MF | 0.36 | 0.004346 | neuro-receptor |
| P2ry6 | MF | 0.35 | 0.006339 | neuro-receptor |
| Lepr | MF | 0.34 | 0.022729 | neuro-receptor |
| Cx3cr1 | MO | 0.54 | 7.47E-11 | cytokine receptor |
| Csf1r | MO | 0.49 | 3.51E-08 | cytokine receptor |

| | | | | |
|---|---|---|---|---|
| Csf3r | MO | 0.47 | 3.97E-07 | cytokine receptor |
| Ltbr | MO | 0.46 | 7.83E-07 | cytokine receptor |
| Tnfrsf1b | MO | 0.43 | 1.24E-05 | cytokine receptor |
| Tnfrsf21 | MO | 0.43 | 1.75E-05 | cytokine receptor |
| Il31ra | MO | 0.40 | 0.000208 | cytokine receptor |
| Il17ra | MO | 0.39 | 0.000286 | cytokine receptor |
| Il13ra1 | MO | 0.39 | 0.000439 | cytokine receptor |
| Tnfrsf1a | MO | 0.38 | 0.001188 | cytokine receptor |
| Csf2ra | MO | 0.37 | 0.002532 | cytokine receptor |
| Il6ra | MO | 0.35 | 0.008121 | cytokine receptor |
| Il10ra | MO | 0.35 | 0.01018 | cytokine receptor |
| Ccr2 | MO | 0.34 | 0.016938 | cytokine receptor |
| Sstr4 | MO | 0.65 | 2.92E-18 | neuro-receptor |
| Adora2b | MO | 0.59 | 8.94E-14 | neuro-receptor |
| S1pr5 | MO | 0.49 | 5.43E-08 | neuro-receptor |
| P2ry6 | MO | 0.47 | 2.05E-07 | neuro-receptor |
| Adrb2 | MO | 0.35 | 0.0078 | neuro-receptor |
| Il18rap | NK | 0.51 | 4.32E-09 | cytokine receptor |
| Il18r1 | NK | 0.48 | 1.19E-07 | cytokine receptor |
| Ccr5 | NK | 0.40 | 0.000229 | cytokine receptor |
| Il2rb | NK | 0.36 | 0.005344 | cytokine receptor |
| Chrne | NK | 0.72 | 2.03E-25 | neuro-receptor |
| S1pr5 | NK | 0.68 | 8.06E-21 | neuro-receptor |
| Hrh2 | NK | 0.38 | 0.001297 | neuro-receptor |
| F2rl2 | NK | 0.36 | 0.003558 | neuro-receptor |
| Cxcr6 | NKT | 0.55 | 3.45E-11 | cytokine receptor |
| Cxcr3 | NKT | 0.48 | 1.52E-07 | cytokine receptor |
| Il18r1 | NKT | 0.42 | 3.10E-05 | cytokine receptor |
| Il12rb1 | NKT | 0.42 | 3.55E-05 | cytokine receptor |
| Tnfrsf25 | NKT | 0.38 | 0.000939 | cytokine receptor |
| Il17rb | NKT | 0.38 | 0.001271 | cytokine receptor |
| Il2rb | NKT | 0.36 | 0.004821 | cytokine receptor |
| Il18rap | NKT | 0.34 | 0.01352 | cytokine receptor |
| Il23r | NKT | 0.34 | 0.018682 | cytokine receptor |
| Hrh4 | NKT | 0.85 | 2.63E-45 | neuro-receptor |
| F2rl2 | NKT | 0.40 | 0.000283 | neuro-receptor |
| F2r | NKT | 0.38 | 0.001129 | neuro-receptor |
| Galr3 | NKT | 0.35 | 0.009889 | neuro-receptor |
| Il17rb | PRET | 0.48 | 6.68E-08 | cytokine receptor |
| Il2ra | PRET | 0.38 | 0.001177 | cytokine receptor |
| Tnfrsf8 | PRET | 0.34 | 0.012123 | cytokine receptor |
| Chrna9 | PRET | 0.53 | 1.80E-10 | neuro-receptor |
| Thra | PRET | 0.44 | 8.81E-06 | neuro-receptor |
| Sstr2 | PRET | 0.42 | 3.97E-05 | neuro-receptor |
| Tnfrsf19 | PROB | 0.74 | 1.63E-27 | cytokine receptor |
| P2rx3 | PROB | 0.85 | 3.04E-45 | neuro-receptor |

| | | | | |
|---|---|---|---|---|
| Mpl | SP | 0.72 | 8.01E-25 | cytokine receptor |
| Il11ra1 | SP | 0.49 | 5.06E-08 | cytokine receptor |
| Il11ra2 | SP | 0.49 | 5.33E-08 | cytokine receptor |
| Csf3r | SP | 0.42 | 2.58E-05 | cytokine receptor |
| Il1r1 | SP | 0.35 | 0.006013 | cytokine receptor |
| Cxcr2 | SP | 0.35 | 0.007022 | cytokine receptor |
| Tnfrsf21 | SP | 0.34 | 0.018463 | cytokine receptor |
| Mc5r | SP | 0.73 | 1.45E-26 | neuro-receptor |
| P2rx1 | SP | 0.57 | 1.09E-12 | neuro-receptor |
| Grik5 | SP | 0.56 | 6.03E-12 | neuro-receptor |
| F2rl3 | SP | 0.56 | 8.95E-12 | neuro-receptor |
| Vipr2 | SP | 0.54 | 7.25E-11 | neuro-receptor |
| Lhcgr | SP | 0.51 | 2.64E-09 | neuro-receptor |
| P2ry2 | SP | 0.48 | 8.43E-08 | neuro-receptor |
| Gabre | SP | 0.47 | 2.29E-07 | neuro-receptor |
| Htr2a | SP | 0.46 | 8.58E-07 | neuro-receptor |
| P2ry1 | SP | 0.42 | 2.45E-05 | neuro-receptor |
| Thra | SP | 0.41 | 6.38E-05 | neuro-receptor |
| Gria3 | SP | 0.41 | 6.64E-05 | neuro-receptor |
| Ghr | SP | 0.35 | 0.00805 | neuro-receptor |
| Cd4 | T4 | 0.65 | 3.42E-18 | cytokine receptor |
| Tnfrsf25 | T4 | 0.53 | 5.25E-10 | cytokine receptor |
| Tnfrsf18 | T4 | 0.41 | 5.69E-05 | cytokine receptor |
| Tnfrsf4 | T4 | 0.41 | 0.000102 | cytokine receptor |
| Il27ra | T4 | 0.38 | 0.000848 | cytokine receptor |
| Tgfbr2 | T4 | 0.38 | 0.001034 | cytokine receptor |
| Ccr7 | T4 | 0.36 | 0.003153 | cytokine receptor |
| Il6st | T4 | 0.35 | 0.007247 | cytokine receptor |
| Il6ra | T4 | 0.35 | 0.00936 | cytokine receptor |
| Cd27 | T4 | 0.33 | 0.033984 | cytokine receptor |
| Gpr83 | T4 | 0.70 | 2.35E-23 | neuro-receptor |
| Vipr1 | T4 | 0.62 | 9.19E-16 | neuro-receptor |
| Gabrr2 | T4 | 0.50 | 7.42E-09 | neuro-receptor |
| S1pr1 | T4 | 0.39 | 0.000323 | neuro-receptor |
| F2rl1 | T4 | 0.36 | 0.003185 | neuro-receptor |
| Il27ra | T8 | 0.34 | 0.015224 | cytokine receptor |
| F2rl1 | T8 | 0.42 | 2.75E-05 | neuro-receptor |
| Gabrr2 | T8 | 0.42 | 4.59E-05 | neuro-receptor |

**Table S8 - Enrichment analysis of NR and cytokine receptors significantly correlated to lineage.** List of enriched Go terms identified for groups of neuro-receptors and cytokine receptors significantly correlated to lineage. For each Go term, we provide information of Benferroni correction p-value, the cytokine receptors and neuro-receptors enriched and the specific lineage.

| Go Term | Adjust pValue | Cytokine receptors | Neuro-receptors | Lineage |
|---|---|---|---|---|
| neutrophil chemotaxis | 0.037 | Csf3r/ Cxcr1/ Cxcr2 | Ednra | GN |
| regulation of cell differentiation | 0.016 | Acvr1/ Ccr1/ Ccr5/ Csf1r/ Csf3r/ Cx3cr1/ Epor/ Tgfbr1/ Tnfrsf12a/ Tnfrsf21 | P2ry2/ Lpar1/ P2rx4/ P2ry12/ Adrb1/ Ednrb | MF |
| neurogenesis | 0.016 | Ccr5/ Csf1r/ Cx3cr1/ Epor/ Tgfbr1/ Tnfrsf12a/ Tnfrsf21 | P2ry2/ Lpar1/ P2rx4/ P2ry12/ Ednrb/ Lepr | MF |
| movement of cell or subcellular component | 0.016 | Acvr1/ Ccr1/ Ccr3/ Ccr5/ Csf1r/ Csf3r/ Cx3cr1/ Tgfbr1/ Tnfrsf12a | P2ry2/ Adora3/ Lpar1/ P2rx4/ P2ry12/ P2ry6/ Adrb1/ Ednrb | MF |
| cell migration | 0.018 | Acvr1/ Ccr1/ Ccr3/ Ccr5/ Csf1r/ Csf3r/ Cx3cr1/ Tgfbr1/ Tnfrsf12a | P2ry2/ Adora3/ Lpar1/ P2rx4/ P2ry12/ P2ry6/ Ednrb | MF |
| negative regulation of angiogenesis | 0.187 | Ccr2/ Cx3cr1 | Adrb2 | MO |
| cytokine production | 0.187 | Ccr2/ Csf1r/ Cx3cr1/ Il17ra/ Il31ra/ Il6ra/ Tnfrsf1a/ Tnfrsf1b/ Tnfrsf21 | Adora2b | MO |
| positive regulation of nitrogen compound metabolic process | 0.187 | Ccr2/ Csf1r/ Cx3cr1/ Il31ra/ Il6ra/ Ltbr/ Tnfrsf1a/ Tnfrsf1b | Adora2b/ P2ry6/ Sstr4/ Adrb2 | MO |
| response to stress | 0.374 | Cxcr3/ Cxcr6/ Il12rb1/ Il17rb/ Il18r1/ Il18rap/ Il23r | F2r/ F2rl2/ Hrh4 | NKT |
| cytokine production | 0.380 | Cxcr3/ Il12rb1/ Il17rb/ Il18r1/ Il18rap/ Il23r | F2r | NKT |
| leukocyte proliferation | 0.003 | Ccr7/ Cd27/ Cd4/ Il6ra/ Il6st/ Tgfbr2/ Tnfrsf4 | F2rl1 | T4 |
| T cell activation | 0.005 | Ccr7/ Cd27/ Cd4/ Il6ra/ Il6st/ Tgfbr2/ Tnfrsf4 | F2rl1 | T4 |
| regulation of cell adhesion | 0.004 | Ccr7/ Cd27/ Cd4/ Il6ra/ Il6st/ Tgfbr2/ Tnfrsf18 | S1pr1 | T4 |
| interleukin-6 secretion | 0.220 | Il27ra | F2rl1 | T8 |

**Table S9 – CyTOF panel for B lymphocytes**

| Protein | Metal | Clone | Extra/intrace cellular | Manufacturer |
|---|---|---|---|---|
| IgMsurface | 115 In | MHM-88 | EC | Biolegend |
| CD19 | 116 Cd | HIB19 | EC | Biolegend |
| CD45* | 141Pr | HI30 | EC | Biolegend |
| CD11c | 142Nd | Bu15 | EC | Biolegend |
| CD117* | 143 Nd | 104D2 | EC | Biolegend |
| CD34* | 144 Nd | 581 | EC | Biolegend |
| CD22 | 145 Nd | HIB22 | EC | Biolegend |
| CD235 | 146 Nd | HIR2 | EC | Biolegend |
| CD20 | 147 Sm | 2H7 | EC | Biolegend |
| CD61 | 148Nd | VI-PL2 | EC | Biolegend |
| CD24 | 149 Sm | ML5 | EC | Biolegend |
| CD123* | 151 Eu | 6H6 | EC | Fluidigm Inc |
| CD16* | 152 Sm | 3G8 | EC | Biolegend |
| CD40 | 153 Eu | 5C3 | EC | Biolegend |
| CD14 | 154Sm | M5E2 | EC | Biolegend |
| CD72 | 155 Gd | 3F3 | EC | Biolegend |
| CD10* | 156 Gd | HI10a | EC | Fluidigm Inc |
| CD179b | 157 Gd | HSL11 | EC | Biolegend |
| CD43 | 158 Gd | 10G7 | EC | Biolegend |
| CD179a | 159 Tb | HSL96 | EC | Biolegend |
| F2R | 160 Gd | 731115 | EC | R&D SYSTEMS |
| GRIA3 | 162Dv | Glu149.29.61 | EC | Novus Biological |
| CD127 | 163 Dy | A019D5 | IC | Biolegend |
| Tdt | 164 Dy | E17-1519 | IC | Biolegend |
| IgM intraecellular | 165Ho | MHM-88 | EC | Biolegend |
| ADRB2 | 166 Er | 586107 | EC | R&D Systems |
| AVPR2 | 168Er | Rbbit polyclonal | EC | Invitrogen |
| CD56 | 169 Tm | HCD56 | EC | Biolegend |
| CD3* | 170 Er | UCHT1 | EC | Biolegend |
| CNR2 | 171Yb | 352110 | EC | R&D Systems |
| CD4 | 173YB | RPA-T4 | EC | Biolegend |
| HLADR | 174 Yb | L243 | EC | Biolegend |

| | | | | |
|---|---|---|---|---|
| CD38* | 175Lu | HIT2 | EC | Fluidigm Inc |
| CD33* | 176 Yb | WM53 | EC | eBioscience |
